# Morphology, Slow Dynamics, and Composition-Controlled Phase Behavior in a Minimal Model of Dps:DNA Co-Condensation

**DOI:** 10.64898/2025.11.28.691232

**Authors:** Alberto Alonso, Soumik Mitra, Luke Studt, Lauren Melcher, Anne S. Meyer, Elio A. Abbondanzieri, Moumita Das

**Affiliations:** School of Mathematics and Statistics, Rochester Institute of Technology, Rochester, NY, USA; School of Physics and Astronomy, Rochester Institute of Technology, Rochester, NY, USA; School of Arts and Sciences, University of Rochester, Rochester, NY, USA

## Abstract

The DNA-binding protein from starved cells (Dps) compacts bacterial DNA into stress-protective condensates, but the physical ingredients that are sufficient to produce the observed morphological and dynamical phenomenology remain unclear. Here we use Dps:DNA as the motivation for a broader soft-matter question: which properties govern the co-condensation of a semiflexible polymer with a binding co-solute, and how is the resulting condensate morphology selected? We combine coarse-grained Brownian dynamics (BD) simulations, in which DNA is a semiflexible bead–spring chain and Dps a spherical particle interacting through heterotypic attraction and homotypic repulsion, with a ternary Flory–Huggins free energy evolved under conserved Cahn–Hilliard–Cook dynamics. Both frameworks show that the condensate morphology is governed jointly by the heterotypic Dps:DNA attraction and by composition: attraction drives co-condensation, while at fixed attraction the relative abundance of the two species selects between extended, network-like structures and compact, droplet-like condensates. That two independent models, sampling different concentration regimes, both show composition-controlled morphology selection indicates it is a robust feature of the co-condensation thermodynamics. The simulations further reveal slow, heterogeneous dynamics: sub-diffusive Dps motion and a stretched-exponential collapse of chain dimensions, signatures of the dynamical arrest that accompanies compaction. The framework provides general, transferable principles for protein–nucleic-acid phase separation in soft and living matter.

## 1 Introduction

The self-organization of DNA and proteins into compartments underlies cellular functions from gene regulation to stress response ^1^. In eukaryotes this organization occurs in membrane-bound organelles and in membrane-less compartments formed by liquid–liquid phase separation (LLPS), where weak multivalent interactions among proteins and nucleic acids create distinct biochemical environments ^2–5^. Bacteria lack internal membranes; their genome is organized within a membrane-less nucleoid that must balance compaction against accessibility, a balance that becomes critical under stress such as starvation or oxidative damage, when many bacteria form stable, membrane-free condensates that compact and protect DNA ^6–9^.

Among bacterial stress proteins, the DNA-binding protein from starved cells (Dps) is a paradigm for multivalent-interaction-driven DNA condensation. Dps is a nucleoid-associated protein expressed at low levels in exponential growth that accumulates dramatically in stationary phase and under stress ^10–14^. Each dodecamer binds DNA through positively charged N-terminal tails, bridging DNA strands into dense assemblies that shield the genome from oxidative and mechanical damage ^15–18^. In vitro, purified Dps rapidly coats DNA to form condensates with liquid-like or crystalline properties ^6^.

DNA and Dps co-condense into liquid-like or other compact morphologies ^6^: a semiflexible polymer and a binding co-solute demixing together. We ask which minimal conditions are sufficient to predict this behavior, and which of them set the condensate morphology. We do not attempt a biochemically detailed model of Dps: we do not resolve its dodecameric structure, tail electrostatics, or sequence dependence, or seek quantitative comparison with specific in vitro measurements. The problem then belongs to the broader class of co-condensation and phase separation, in which a binding partner drives demixing of a polymer that would not phase-separate on its own: Dps:DNA serves as the motivating example, and the resulting physical principles apply more generally.

We use two complementary models. Coarse-grained BD simulations resolve the microscopic organization and dynamics of the co-condensate, including how interaction strength and stoichiometry produce liquid-like or extended morphologies. A ternary Flory–Huggins free energy evolved under conserved Cahn–Hilliard–Cook dynamics provides the continuum description: it locates the thermodynamic regimes of demixing and follows the pathway by which a homogeneous mixture organizes into network-like extended structures or droplet-like condensates. The two models resolve different physics, so we compare them at the level of morphological phase behavior rather than parameter by parameter. Together they link microscale interactions and composition to the mesoscale organization of the condensate.

## 2 Model and Methods

### 2.1 Brownian Dynamics Simulations

We investigated the microscopic organization and dynamics of Dps:DNA mixtures using quasi 2D coarse-grained BD simulations, which provide microscale insights into condensate morphology and dynamical behavior. DNA was modeled as polymer chains of 500 beads connected by harmonic springs, each bead having an effective diameter of 3 nm (experimentally closer to 2 nm). The 3 nm value was selected to achieve DNA chain lengths comparable to experimentally relevant contour lengths. Dps proteins were modeled as 9 nm spherical colloids ^10^, approximately three times the diameter of a DNA bead. Systems containing 100–3000 Dps particles and 3–9 DNA chains were simulated over a range of stoichiometries and interaction strengths, with a hard-wall repulsive boundary for the simulation box to prevent DNA wrapping and the solvent treated implicitly. The simulation box had dimensions 200*σ* × 200*σ* × 15*σ*, consistent with quasi 2D geometry. The quasi-two-dimensional geometry was chosen to approximate the geometry of in-vitro experiments ^19^ and to keep the accessible system size tractable for simulations with explicit semiflexible chains; as shown below, the continuum model solved in a periodic two-dimensional bulk reaches the same morphology classes, indicating that the observed morphologies are not artifacts of the slab confinement.

Heterotypic Dps:DNA interactions were modeled with a Lennard–Jones potential that captures both excluded-volume repulsion and medium-range attraction, with the attraction strength tuned to mimic the effects of salt concentration and macromolecular crowding (e.g. PEG) on Dps:DNA binding affinity. The LJ potential is defined as

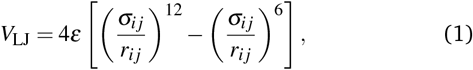

where *r*_*i j*_ is the center-to-center distance between beads *i* and *j, ε* denotes the interaction strength, and *σ*_*i j*_ is the arithmetic mean of the bead diameters. In contrast, homotypic interactions, namely DNA:DNA and Dps:Dps, were modeled using the purely repulsive Weeks–Chandler–Andersen (WCA) potential, described by *V*_WCA_(*r*) = *V*_LJ_(*r*), *r <* 2^1*/*6^*σ*, and *V*_WCA_(*r*) = 0 otherwise which ensures excluded-volume effects without any attractive tail, thereby preventing unphysical aggregation. Together, this interaction scheme allows Dps and DNA to form condensates via heterotypic attraction, while maintaining steric stability through homotypic repulsions.

The motion of each DNA bead *a* was governed by an over-damped Langevin equation:

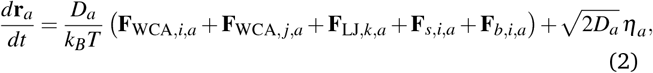

where *D*_*a*_ is the diffusion constant for the DNA beads, *k*_*B*_ is the Boltzmann constant, *T* is the temperature, and *η*_*a*_ is a Gaussian random force introducing thermal noise. The forces **F**_WCA,*i,a*_, **F**_WCA, *j,a*_, and **F**_LJ,*k,a*_ represent pairwise interaction forces acting on bead *a*. Index *i* corresponds to intra-strand interactions between beads on the same DNA strand, index *j* to inter-strand DNA–DNA interactions, and index *k* to interactions between DNA beads and Dps particles. Bonded DNA beads within a chain experience stretching and bending forces to preserve chain elasticity. The stretching energy is given by 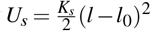, and the bending energy by 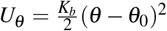. Here, *K*_*s*_ is the extensional stiffness coefficient, *l* is the distance between consecutive beads along a strand, and *l*_0_ is the equilibrium bond length, set equal to the bead diameter. The bending stiffness is represented by *K*_*b*_, where *θ* is the angle between three consecutive beads and the equilibrium angle *θ*_0_ is set to *π*. All forces were obtained by calculating the appropriate negative gradients of the interaction or mechanical energies. The translational motion of each Dps particle *b* was described by a similar overdamped Langevin equation:

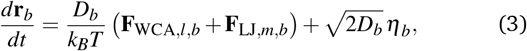

where *l* denotes Dps:Dps interactions and *m* represents Dps:DNA interactions, *D*_*b*_ is the diffusion constant for the Dps beads, and *η*_*b*_ is a Gaussian random force introducing thermal noise.

The equations were non-dimensionalized as follows. All lengths were non-dimensionalized by the DNA bead diameter *σ* = 3 nm, and all times by *t*_0_ = 0.3 *µ*s corresponding to the diffusion time of a single DNA bead across its own diameter, with the diffusion coefficient *D*_*a*_ estimated from prior work ^20,21^. Forces were non-dimensionalized by *k*_*B*_*T/σ*. DNA chains were first equilibrated for *t* = 1000*t*_0_ before introducing Dps particles; each simulation then ran for up to *t* = 14000*t*_0_. Rather than fixing a stopping time, we assessed convergence from time-resolved order parameters. The principal observable is a segment-averaged radius of gyration: each 500-bead chain is partitioned into consecutive 100-bead segments, the *R*_*g*_ of each segment is computed and averaged over the segments of a chain to give one value per chain, and these per-chain values are averaged over the chains in the system and over four independent seeds. The SEM is the sample standard deviation of the per-chain values divided by 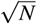, with *N* = *N*_chains_ × *N*_seeds_; segments are averaged within a chain rather than treated as independent samples. This segment-averaged *R*_*g*_(*t*) decreases from its initial extended value as the chains compact and reaches a plateau at late times, with the mean ± standard-error of mean (SEM) band narrowing and the plateau reproducible across seeds. The band can be relatively wide because the 100-bead segmentation samples a heterogeneous population of conformations, with some segments in elongated arms and others in compact cores, so individual chains span a broad range of *R*_*g*_ even where the mean is well determined. Because *R*_*g*_(*t*) plateaus while condensate domains continue to coarsen slowly, the appropriate description is a statistical steady state with slow, arrested coarsening rather than equilibration to a single condensate. The configurations shown are representative of the morphology class reproduced across seeds.

### 2.2 Thermodynamic model for phase behavior

We employed a Flory–Huggins (FH) free-energy framework ^22–24^ to model the thermodynamics of the ternary Dps:DNA:solvent mixture. In this coarse-grained lattice model, polymers and small molecules occupy discrete sites and interact via effective pairwise potentials, capturing the balance of entropic mixing and enthalpic interactions that drives phase separation. The FH theory identifies compositions that favor Dps:DNA condensate formation; the BD simulations discussed above then provide structural and kinetic detail that complements these thermodynamic predictions. For concentrations *ϕ*_DNA_, *ϕ*_Dps_, and *ϕ*_Solv_, the FH free energy is

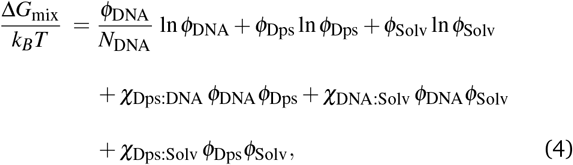

where *k*_*B*_ is the Boltzmann constant, *T* is temperature, and *N*_DNA_(~ 10^3^) is a measure of DNA polymerization. Further, we use volume fractions as our measure of concentration. The *χ*_*i j*_ are FH interaction parameters. For nearest–neighbor coordination number and pair interaction energies 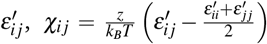. For the ternary mixture these become, 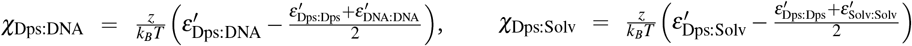, and 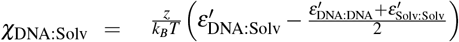 A negative *χ*_Dps:DNA_ corresponds to energetically favorable heterotypic attraction between Dps and DNA, promoting their mutual association, whereas positive *χ*_DNA:Solv_ and *χ*_Dps:Solv_ penalize solute–solvent contacts, thus promoting phase separation between Dps:DNA-rich and solvent-rich phases. Notably, the pair interaction energies 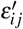 entering the Flory–Huggins free energy are effective coarse-grained thermodynamic interaction parameters and should not be interpreted as the microscopic Lennard– Jones interaction strengths *ε*_*i j*_ used in the Brownian dynamics simulations. The two also follow different sign conventions: in the Brownian dynamics simulations *ε* is a Lennard–Jones well depth, so a larger positive *ε*_Dps:DNA_ denotes stronger interaction, whereas the Flory–Huggins pair energies 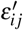 and the resulting *χ*_*ij*_ follow the convention described above. Accordingly, neither 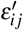 nor the resulting Flory–Huggins parameters *χ*_*ij*_ are intended to map quantitatively onto the Brownian dynamics interaction parameters; rather, they provide an effective thermodynamic description of the same qualitative co-condensation behavior.

We evaluated Eq (4) over the incompressible simplex (*ϕ*_DNA_ + *ϕ*_Dps_ + *ϕ*_Solv_ = 1), identified spinodal regions from the Hessian of the free-energy density, and obtained binodal boundaries from the coexistence conditions. These maps elucidate how Dps:DNA attraction and effective solute–solvent repulsions/solvent quality cooperatively control condensate formation. Two phases coexist at equilibrium when the chemical potentials of each independent component and the osmotic pressures are equal in both phases, and the total free energy is minimized.

Because the mixture is incompressible, *ϕ*_Solv_ = 1 − *ϕ*_DNA_ − *ϕ*_Dps_, so only two compositions are independent. We eliminate the solvent and treat it as the reference species, writing the free-energy density as a function of the two independent fields,

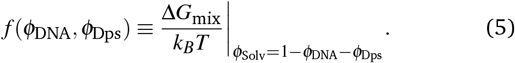

The exchange chemical potentials of the independent species follow as

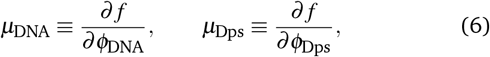

each taken at fixed value of the other independent field; the derivative with respect to the solvent fraction does not appear, as the solvent is fixed by incompressibility. Two phases (labels 1 and 2) coexist when these exchange potentials and the osmotic pressure are equal in both phases,

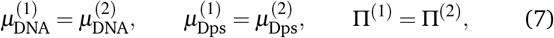

where the osmotic pressure is Π = *µ*_DNA_*ϕ*_DNA_ + *µ*_Dps_*ϕ*_Dps_ − *f*.

### 2.3 Conserved Cahn–Hilliard–Cook dynamics

To follow the dynamical pathway of condensate formation, and not only the equilibrium boundaries, we evolve the ternary mixture under conserved Cahn–Hilliard–Cook dynamics. Interfacial penalties are added to the free energy, in units of *k*_*B*_*T*, through a square-gradient functional,

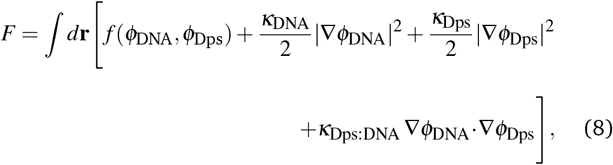

where the *κ* set the interfacial widths. The two independent composition fields evolve as

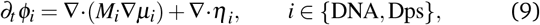

with mobilities *M*_*i*_, chemical potentials *µ*_*i*_ = *δ F/δϕ*_*i*_ that combine the local Flory–Huggins term with gradient contributions, and a weak, conserved stochastic flux noise *η*_*i*_ obeying the fluctuation–dissipation relation. The solvent field follows from incompressibility, so the total amounts of DNA and Dps are conserved. The homogeneous state is linearly unstable (spinodal) where the Hessian of *f* (*ϕ*_DNA_, *ϕ*_Dps_) acquires a negative eigen-value; deep quenches inside the spinodal coarsen into bicontinuous structures, while compositions near the boundary phase-separate by nucleation into droplets. The equations are solved on a periodic two-dimensional domain by a Fourier pseudospectral method with semi-implicit treatment of the gradient terms; full implementation details are given in the SI.

## 3 Results and Discussion

### 3.1 Morphological transitions and condensate formation

Brownian dynamics (BD) simulations reveal that condensate morphology is highly sensitive to the heterotypic Dps:DNA attraction and the Dps concentration. Figure 1 shows steady-state snapshots across a range of attractive interaction strengths (columns) and Dps concentrations (rows). At low attraction and low *ϕ*_Dps_, DNA remains in extended, network-like configurations with dispersed Dps. Increasing the interaction strength produces branched aggregates, and at the strongest attraction and moderate *ϕ*_Dps_, compact condensates with dense Dps:DNA cores and dilute surroundings. Within each column, higher *ϕ*_Dps_ drives a similar transition to more tightly condensed morphologies; thus Fig. 1 shows that both axes matter, but in distinct ways: the heterotypic attraction controls whether dense clusters form at all, whereas at fixed attraction the composition sets which morphology, branched network or compact droplet, is selected. We consider a Dps particle to be a part of the condensate if it is within ~ 6 nm of another Dps bead or a bead in the DNA chain which is a part of the condensate; this threshold is the Dps:DNA contact distance *σ*_*i j*_ = (9 + 3)*/*2 nm, i.e. the first-neighbor shell, so a particle is counted as bound once it is within one interaction range of the condensate.

Figure 2 complements this view by mapping morphologies across Dps and DNA concentrations at a fixed attraction (*ε*_Dps:DNA_ = 1.5). In the absence of Dps (*ϕ*_Dps_ = 0, left column), the DNA remains in extended, self-avoiding coils and does not condense. At fixed *ϕ*_DNA_, increasing *ϕ*_Dps_ draws the DNA into compact, isolated droplets. At the lowest *ϕ*_DNA_ this progression saturates: once the available DNA is coated, the excess Dps remains dispersed as a dilute, unbound structure (bottom right), so condensation is limited by the amount of DNA rather than by the attraction strength. At fixed *ϕ*_Dps_, raising *ϕ*_DNA_ increases connectivity, taking the system from isolated globular droplets through bridged aggregates to percolated, network-like structures that span the box at the highest DNA loading. Throughout, the phase map highlights local heterogeneity, with dense Dps:DNA condensates coexisting with dilute regions. Overall, the rows and columns indicate that the compact, droplet-like morphology occupies an intermediate window: within the sampled range, a relative excess of DNA drives extended, percolated networks, whereas an excess of Dps beyond what the available DNA can bind leaves the surplus dispersed as an unbound gas. The two routes out of the droplet window are thus physically distinct: added DNA scaffolds connectivity, while added Dps saturates and disperses, rather than a single symmetric trend. Together, Figs. 1 and 2 show how the Dps:DNA interaction strength and the Dps:DNA stoichiometry govern the emergence and morphology of Dps:DNA condensates.

**Fig. 1.**
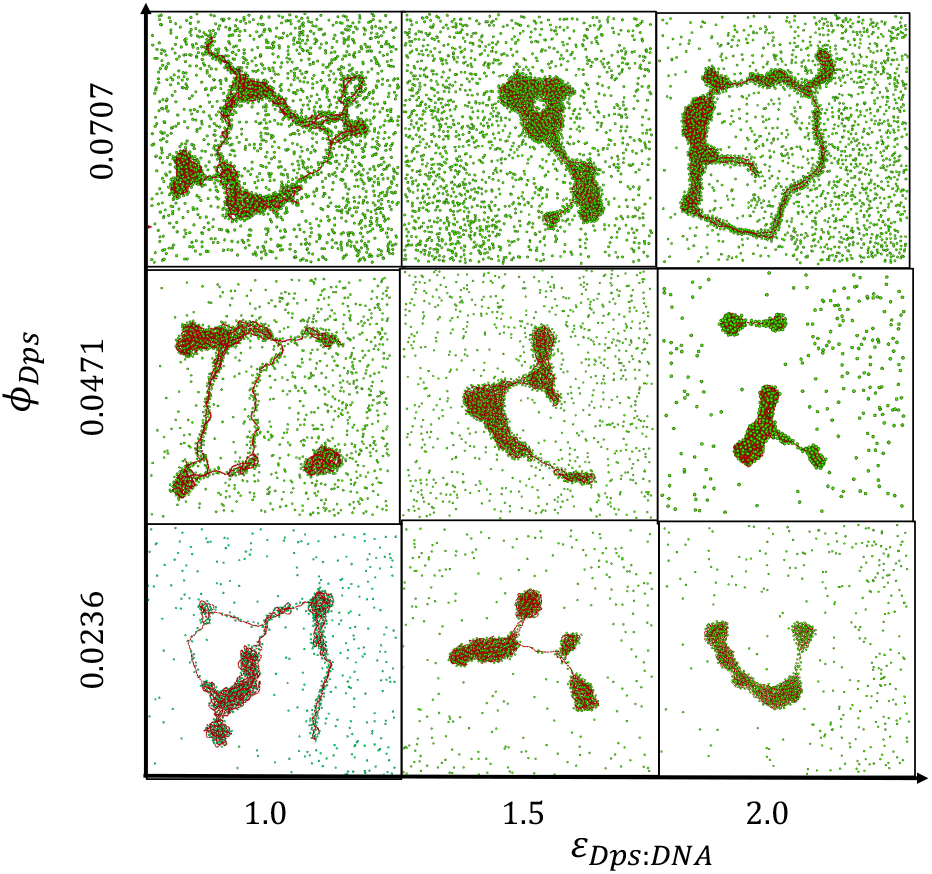
Simulation configurations as a function of Dps concentrations, *ϕ*_*Dps*_ (y-axis), and Dps:DNA interaction strengths, *ε*_*Dps*:*DNA*_ (x-axis). The DNA concentration is fixed at *ϕ*_*DNA*_ = 0.0026, and the Dps:Dps and DNA:DNA interaction strengths are fixed at *ε*_*Dps*:*Dps*_ = *ε*_*DNA*:*DNA*_ = 1. Increasing *ϕ*_*Dps*_ and *ε*_*Dps*:*DNA*_ promotes the formation of compact condensates and alters the phase boundaries of the system.

**Fig. 2.**
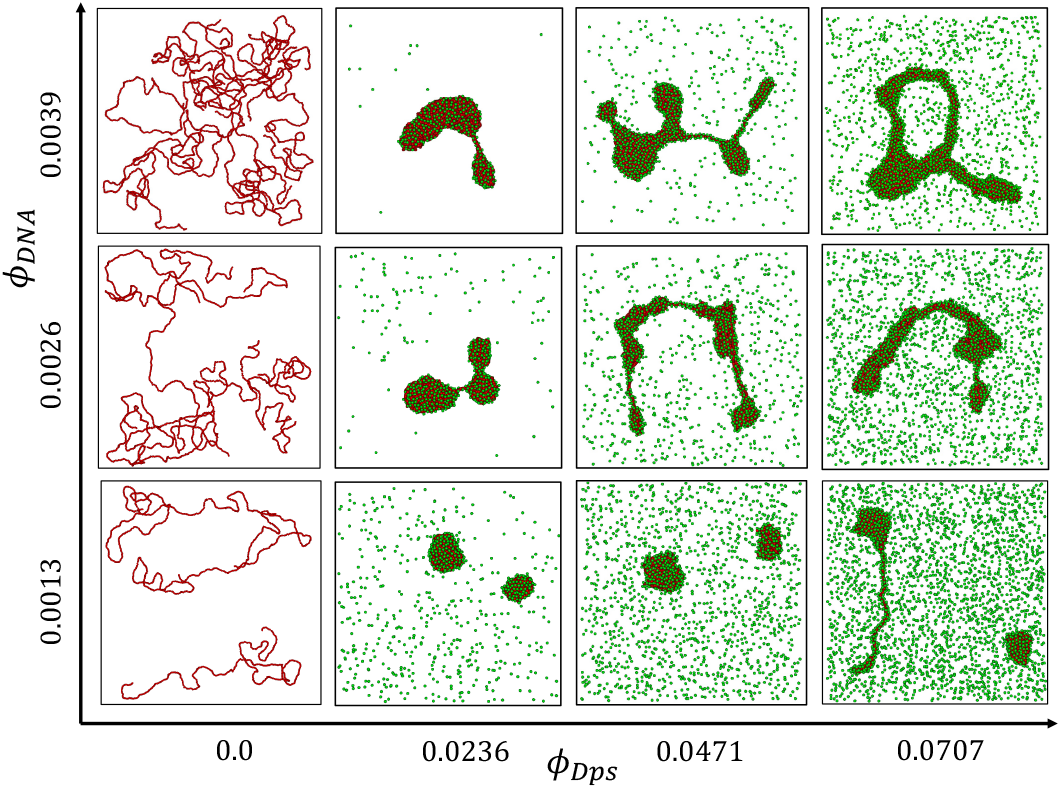
Simulation configurations as a function of Dps concentrations, *ϕ*_*Dps*_ (x-axis), and DNA concentrations,*ϕ*_*DNA*_ (y-axis). The interaction strength between Dps and DNA was kept fixed at *ε*_*Dps*:*DNA*_ = 1.5. The DNA strands are visualized first (in red) and the Dps particles are drawn in subsequently (in green). Volume fraction values of *ϕ*_*Dps*_ = 0.0236, 0.0471, 0.0707 correspond to Dps particle values of 1000, 2000, and 3000, respectively, and volume fraction values of *ϕ*_*DNA*_ = 0.0013, 0.0026, 0.0039 correspond to 3, 6, and 9 chains of DNA.

### 3.2 Dps dynamics and sub-diffusive motion within condensates

We quantified Dps mobility by computing 2D (x–y) mean-squared displacements (MSDs) across a range of DNA and Dps concentrations, *ϕ*_DNA_ and *ϕ*_Dps_ (Fig. 3). For each condition, we plotted MSD versus lag time *τ* for three classes of particles: Dps that ultimately become incorporated into condensates (“eventually bound,” red), Dps that remain outside condensates (“free,” blue), and the ideal free-diffusion reference 4*Dτ* (magenta). All curves are shown on log–log axes.

**Fig. 3.**
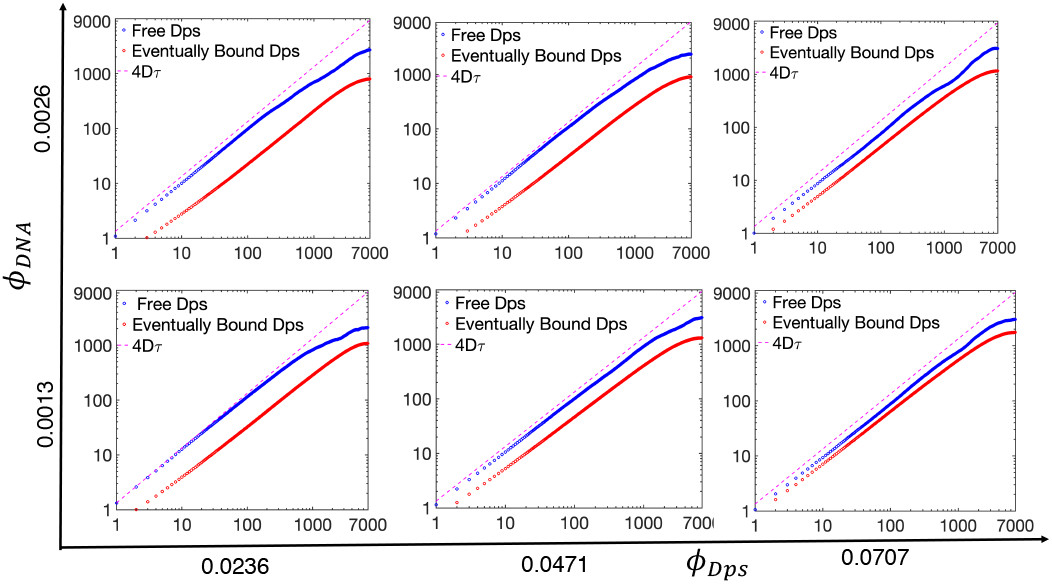
MSD vs Lag Time, *τ*, for different concentrations of Dps (x-axis) and DNA (y-axis). The interaction parameters are *ε*_*Dps*:*Dps*_ = 1, *ε*_*DNA*:*DNA*_ = 1 and *ε*_*Dps*:*DNA*_ = 1.5. The MSD in this diagram is computed for Dps particles that become a part of a condensate (red) and those that do not (blue). The magenta line 4*Dτ* denotes the free diffusion reference (slope 1 on log-log axes); a smaller slope indicates sub-diffusive motion.

Across all parameter combinations, both free and eventually bound Dps exhibit diffusive behavior at short to intermediate lag times (*τ* ≲ 10^2^*t*_0_). The free Dps diffuse slightly more slowly than the 4*Dτ* benchmark, consistent with weak hindrance due to repulsive interactions with other Dps or confinement effects. The eventually bound population shows an even lower effective diffusion constant at these early times, suggesting that these particles begin to experience local crowding or nascent clustering well before fully entering a condensate.

At longer lag times (typically a few ×10^3^*t*_0_), both classes deviate from simple diffusion and enter a sub-diffusive regime. This crossover indicates the onset of increasingly constrained motion, arising from transient interactions, emergent network-like or extended structures, or the approach to condensate formation.

Single-particle tracking further clarifies this transition. Figure 4(a) shows the trajectory of a representative Dps particle in the *xy* plane, with time encoded by color. Early in the simulation, the particle performs a random walk with relatively large displacements (cool colors); after encountering a DNA strand and nearby Dps particles, its motion becomes confined to a much smaller region (warm colors), consistent with the reduced MSD observed in Fig. 3 for large lag times (*τ* ≳ 10^3^*t*_0_).

**Fig. 4.**
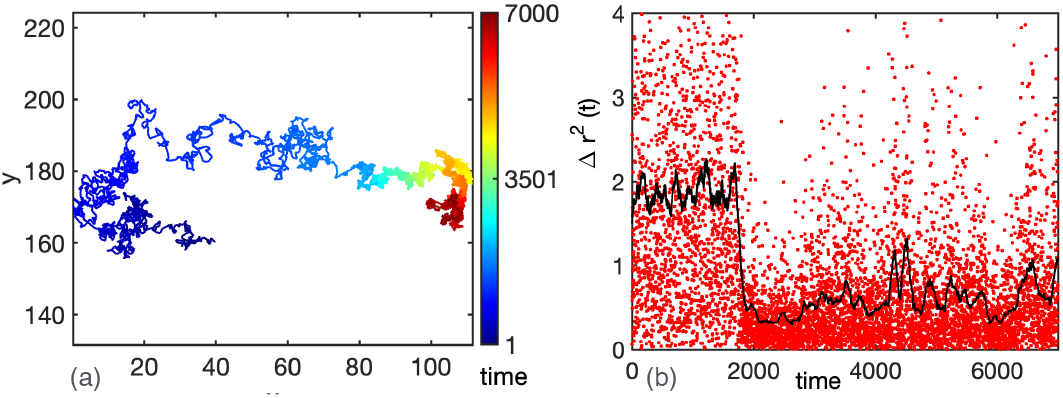
Single particle tracking of a tagged Dps particle that becomes part of a cluster for the case where *ϕ*_*DNA*_ = 0.0026 and *ϕ*_*Dps*_ = 0.0471. (*a*) The particle starts off in a random walk-like motion before its movement becomes restricted due to interactions with DNA and other Dps particles. We can see the particle’s diffusive movement slows down for *t >* 2000, denoted in the warmer-toned colors on the heat map. The heatmap here indicates time. (*b*) Shows the square of the particle’s displacement per time step, and a sliding average of every 100 data points, as a function of time.

This transition is seen even more clearly in Fig. 4(b), where we show the squared displacement per timestep together with a sliding 100-point average. This reveals a sharp decrease in the diffusive movement just before 2000 *t*_0_, marking the onset of confinement. These results illustrate how a Dps particle can abruptly transition from a freely diffusing state to being trapped within a condensate, drastically slowing its dynamics. Together, the MSD analysis and single-particle tracking show a clear link between morphological compaction and dynamic arrest in the Dps:DNA system.

Beyond single-particle motion, the collapse of the chains themselves is slow and heterogeneous. The segment-averaged radius of gyration *R*_*g*_(*t*) (Fig. 5) decreases and plateaus, and its time dependence is well described by a stretched exponential 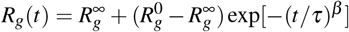, with *β* ≈ 0.6 across the Dps and DNA concentrations studied. A stretching exponent below unity reflects a broad, heterogeneous spectrum of relaxation times, consistent with crowded, glassy compaction rather than a single-rate collapse. This stretched-exponential chain collapse is distinct from the coarsening of condensate domains in the continuum thermodynamic theory.

**Fig. 5.**
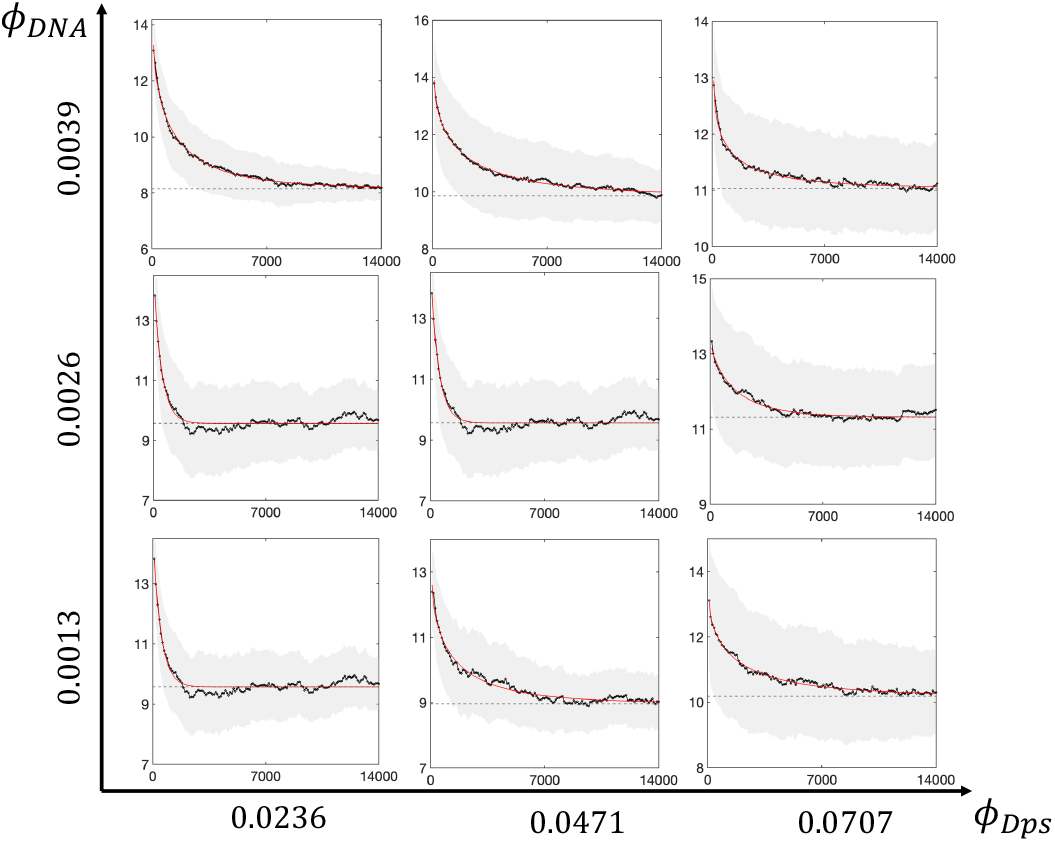
DNA chain compaction and approach to a statistical steady state. Segment-averaged radius of gyration 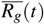 for DNA concentrations *ϕ*_DNA_ (rows, top to bottom: 0.0039, 0.0026, 0.0013) and Dps concentrations *ϕ*_Dps_ (columns, left to right: 0.0236, 0.0471, 0.0707). Black: mean; grey band: ± one standard error of mean (SEM); red: stretched-exponential fit. In every case 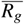 decreases from its extended initial value and plateaus, indicating chain compaction and a statistical steady state. The stretching exponent *β* ≈ 0.6 reflects a broad spectrum of relaxation times characteristic of crowded, arrested collapse. Interaction strengths: *ε*_Dps:DNA_ =1.5, *ε*_Dps:Dps_ = *ε*_DNA:DNA_ =1.

### 3.3 Thermodynamic phase behavior and comparison with simulations

To elucidate the thermodynamic basis of condensate formation, we calculated Flory–Huggins (FH) free-energy landscapes for the ternary Dps:DNA:solvent mixture across a wide range of component concentrations and interaction parameters. The total free energy combines entropic contributions that favor mixing and enthalpic terms that stabilize or destabilize the mixture depending on the sign and magnitude of the Flory–Huggins interaction parameters *χ*_*i j*_. By varying the DNA and Dps concentrations, *ϕ*_DNA_ and *ϕ*_Dps_, from 0.001 to 0.999 (subject to *ϕ*_Solv_ = 1 − *ϕ*_DNA_ − *ϕ*_Dps_), we obtained a free-energy surface *f* (*ϕ*_DNA_, *ϕ*_Dps_) shown as a heat map in Fig. 6.

**Fig. 6.**
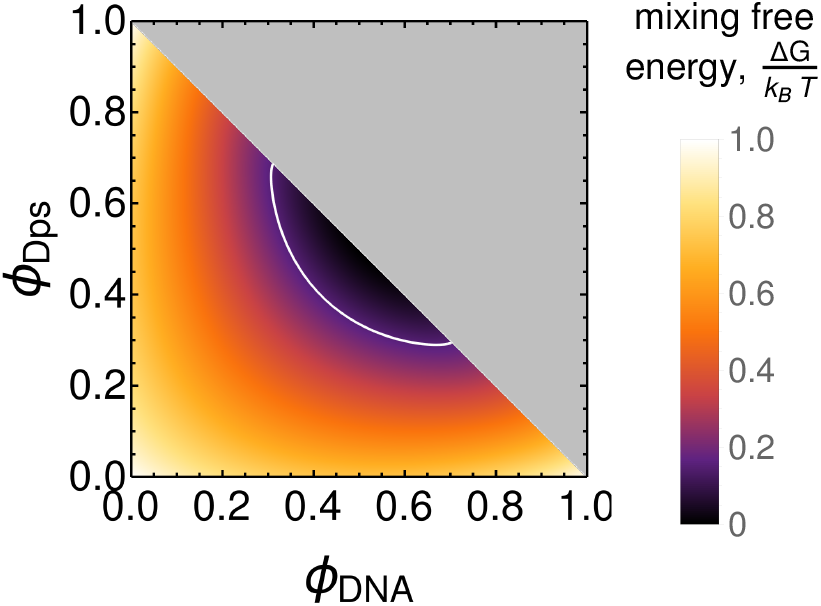
Phase diagram of the ternary Dps:DNA:solvent mixture, projected onto the Dps–DNA composition plane (the solvent fraction is fixed by incompressibility, *ϕ*_Solv_ = 1 − *ϕ*_DNA_ − *ϕ*_Dps_). Flory–Huggins parameters used: *χ*_Dps:DNA_ = −1.25, *χ*_DNA:Solv_ = 1.50, *χ*_Dps:Solv_ = 1.25. The composition of Dps and DNA along the bright (binodal) contour corresponds to the coexistence phases in stable equilibria. The area enclosed by the binodal contour identifies the condensate formation regime for the interacting Dps and DNA molecules. The negative *χ*_Dps:DNA_ together with positive solute–solvent interaction parameters promotes Dps:DNA association while driving demixing from the solvent, thereby enabling condensate formation. The gray triangular region is compositionally inaccessible under the incompressibility constraint. The plotted mixing free energy is rescaled to lie between 0 and 1.

Dark regions correspond to thermodynamically stable homogeneous mixtures, whereas lighter regions indicate higher free-energy states. The white contour denotes the binodal separating the single-phase and two-phase regions of the phase diagram. Compositions inside the binodal undergo phase separation into a dense Dps–DNA-rich condensate coexisting with a solvent-rich dilute phase, while those outside remain homogeneous. The coexistence region is widest for comparable Dps and DNA concentrations, where favorable heterotypic interactions are most effectively realized, and shrinks when either component is present in large excess. This behavior is consistent with the negative Dps–DNA interaction parameter together with unfavorable solute–solvent interactions, which collectively drive co-condensation. The phase diagram thus provides the thermodynamic basis for the morphology selection explored in the subsequent Cahn–Hilliard simulations.

Figure 7 illustrates the emergence of distinct condensate topologies in the ternary Dps–DNA–solvent system as a function of mean composition. Starting from an initially homogeneous state, the composition field undergoes spontaneous phase separation. At an excess of Dps over DNA, the mixture evolves into isolated droplet-like condensates through a nucleation-and-growth-like pathway. As the DNA fraction is raised toward parity with Dps, the morphology crosses over to an interconnected, network-like morphology characteristic of spinodal decomposition. The temporal evolution further reveals the coarsening dynamics of both morphologies, with droplet states growing through coalescence and ripening, whereas bicontinuous domains coarsen by domain thickening and merging. These results highlight that, in addition to interaction strengths, the relative stoichiometry of DNA and Dps plays a central role in selecting the final condensate architecture.

**Fig. 7.**
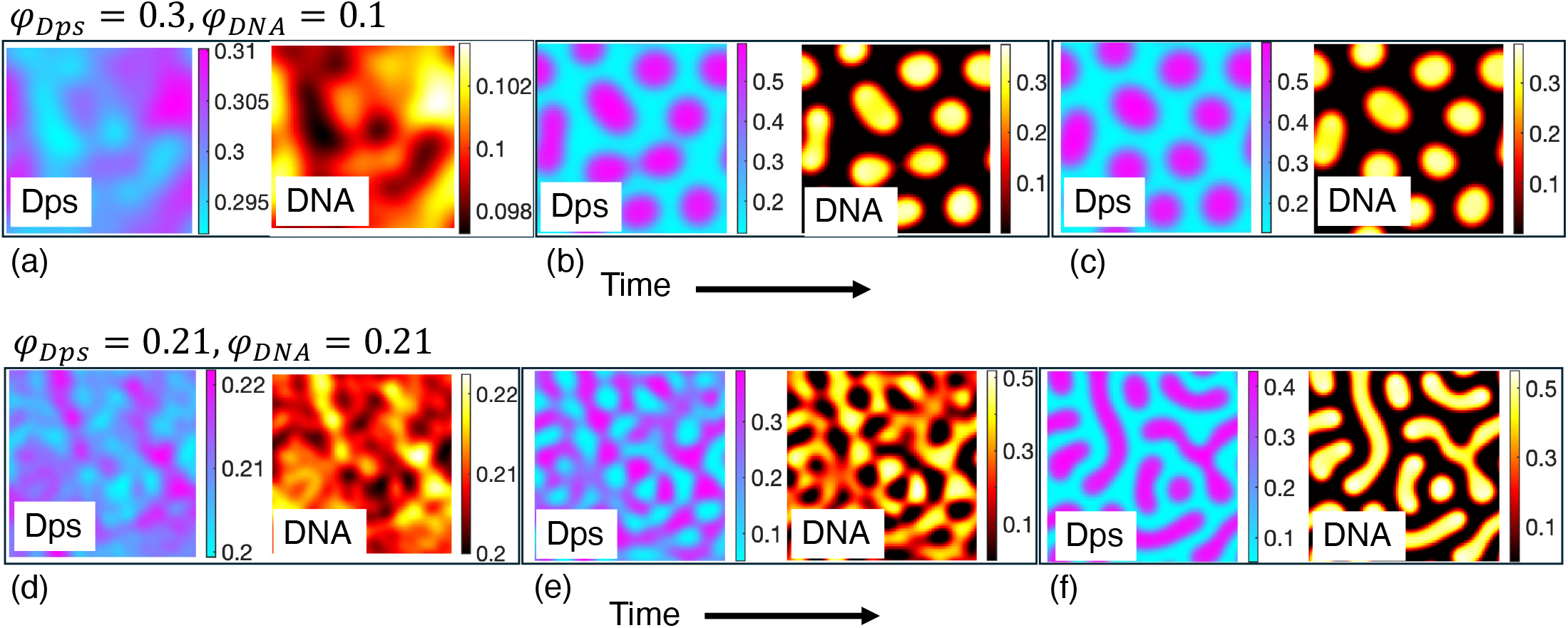
Morphological evolution in a ternary Dps:DNA:solvent system described by a Flory–Huggins–Cahn–Hilliard–Cook model. Each panel shows the Dps volume fraction *ϕ*_Dps_ (left) and the DNA volume fraction *ϕ*_DNA_ (right) at the same instant, plotted with separate colormaps and independent colorbars. The colorbars are scaled to each field and each time point, so levels are read from the individual colorbars and are not comparable across panels; Time increases from left to right within each row. Top row (a-c): Mean composition (*ϕ*_Dps_, *ϕ*_DNA_) = (0.3, 0.1), with Dps in excess of DNA, which coarsens into isolated globular droplets. Bottom row (d-f): Mean composition (*ϕ*_Dps_, *ϕ*_DNA_) = (0.21, 0.21), which percolates into an interconnected, bicontinuous Dps:DNA-rich morphology coexisting with a solvent-rich phase. Thus, at much less DNA than Dps the system forms droplets, while increasing the DNA volume fraction promotes elongated, extended or network-like condensates, consistent with DNA acting as the connectivity-providing scaffold (cf. Fig. 2). Flory-Huggins parameters used: *χ*_Dps:DNA_ =− 2.0, *χ*_Dps:Solv_ = 2.0, *χ*_DNA:Solv_ = 2.0. The interfacial penalties for droplets (a-c): *κ*_Dps:DNA_ = 1.8, *κ*_Dps_ = *κ*_DNA_ = 2.4; network-like (d-f): *κ*_Dps:DNA_ = 0.85, *κ*_Dps_ = *κ*_DNA_ = 1.1. Equal mobilities were assumed for all species. The fields were evolved on a two-dimensional 128 × 128 grid.

The conserved Cahn–Hilliard–Cook dynamics access the same morphology classes as the BD simulations under composition control (Fig. 8). The two descriptions sample different regions of composition space: the BD state points sit in the dilute limit and with large Dps-to-DNA excess (*ϕ*_Dps_*/ϕ*_DNA_ ≈ 6–54), whereas the Cahn–Hilliard maps are evaluated near balanced composition of Dps and DNA and comparatively high total volume fractions of the two. In BD the solvent is implicit, so only *ϕ*_Dps_ and *ϕ*_DNA_ are controlled, and DNA is represented explicitly as a semiflexible chain, so its mechanical properties influence the morphology directly. The composition-field Cahn–Hilliard–Cook model instead evolves *ϕ*_Dps_, *ϕ*_DNA_, and *ϕ*_Solv_ as conserved fields coupled through Flory–Huggins interactions, and carries no explicit polymer degrees of freedom: chain connectivity and semiflexibility are not represented. Because Dps and DNA co-localize in a single dense phase in both descriptions, we compare them through the morphology of the resulting co-condensate: in the composition-field model, through the spatial organization and co-localization of the *ϕ*_Dps_ and *ϕ*_DNA_ fields, and in the BD simulations through the shape of the mixed Dps:DNA-rich domains. We therefore compare them only at the level of morphology class. That two such different schemes nevertheless select the same morphologies indicates the behavior reflects the co-condensation thermodynamics common to both, rather than the details of either.

**Fig. 8.**
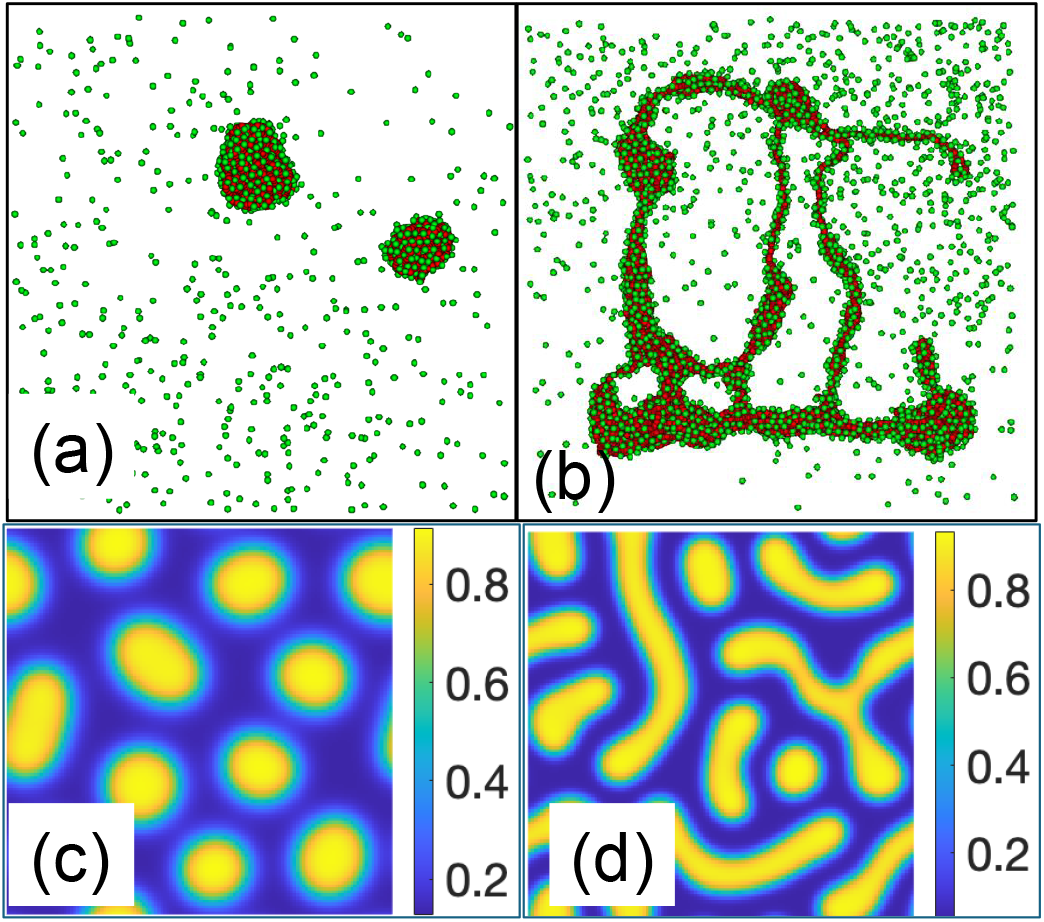
Composition-controlled morphology of Dps:DNA co-condensates in two independent coarse-grained descriptions. In both descriptions Dps and DNA co-condense into a shared dense phase, and we characterize the morphology of that co-condensate rather than of either component alone. Top row, Brownian Dynamics (BD) (a)–(b): the Dps (green) and DNA (red) particles occupy the same dense regions, forming isolated compact co-condensates (a) and an extended, network-like co-condensate (b). Bottom row, Flory–Huggins–Cahn–Hilliard–Cook (CHC) (c)–(d): the color encodes the total co-condensate composition *ϕ*_Dps_ + *ϕ*_DNA_ (shared colorbar; dense co-condensate in yellow, solvent-rich background in blue), giving droplet (c) and extended (d) morphologies. In both descriptions, an excess of Dps over DNA yields isolated droplets, whereas increasing the DNA fraction drives elongated, network-like co-condensates. Absolute concentrations are not directly comparable between the two descriptions; the comparison is made at the level of these qualitative morphological trends.

## 4 Conclusions

We developed a minimal framework combining coarse-grained Brownian dynamics with a ternary Flory–Huggins free energy evolved under conserved Cahn–Hilliard–Cook dynamics, using Dps:DNA as the motivating example of polymer-co-solute co-condensation. In the dilute regime probed by the simulations, both the Dps:DNA attraction and the composition shape condensate morphology: attraction drives co-condensation, and at fixed attraction the stoichiometry selects the morphology. With Dps in large excess over DNA throughout, compact droplet-like condensates form in a narrow window at low DNA content, and increasing the DNA fraction drives the crossover to elongated or network-like structures; the surplus Dps that the available DNA cannot bind remains dispersed as an unbound gas. This compaction is accompanied by dynamical arrest. Dps incorporated into condensates becomes effectively trapped and moves sub-diffusively, while unbound Dps remains nearly Brownian, and the chains themselves collapse with a time course well described by a stretched exponential (*β* ≈ 0.6), indicating a broad spectrum of relaxation times rather than a single-rate process.

The continuum model provides the complementary thermodynamic and kinetic picture at higher overall concentrations. For the interaction parameters considered here, the Flory–Huggins free energy indicates that co-condensation is most favorable near balanced Dps:DNA compositions and is suppressed when either component is present in large excess, and the Cahn–Hilliard– Cook dynamics resolve the kinetic pathways to the resulting morphologies: spinodal decomposition into connected, bicontinuous structures near balance, and nucleation and growth of isolated droplets under strong stoichiometric asymmetry, consistent with established phase-field treatments of multicomponent mixtures ^25–29^. Together, the two approaches indicate that condensate morphology is set jointly by interactions and composition, with composition selecting between connected and isolated morphologies at fixed interactions.

The agreement between the two descriptions, despite their different microscopic physics and non-overlapping composition ranges, indicates that composition-controlled morphology selection is set by the co-condensation thermodynamics rather than by the details of either model. Neither model resolves Dps dodecameric structure, tail electrostatics, or sequence-dependent DNA mechanics, so we frame the contribution as general physical principles for protein–nucleic-acid co-condensation. Quantitative comparison with specific in vitro observables is a natural next step, as are extensions to electrostatic screening, multivalent ions, charge heterogeneity, and sequence-dependent flexibility, and application to other nucleoid-associated proteins and chromatin-like systems.

## Author contributions

Conceptualization: M.D. Methodology: A.A., S.M., L.S., L.M., M.D. Investigation: A.A., S.M., L.S., L.M., M.D. Visualization: A.A., S.M., L.S., L.M., E.A., M.D. Formal Analysis: A.A., S.M., E.A., A.S.M., M.D. Supervision: M.D. Writing, original draft: A.A., S.M., M.D. Writing, review & editing: A.A., S.M., E.A., A.S.M., M.D.

## Data availability

The simulation and analysis code, together with the data underlying the figures in this article, are available from the corresponding author upon publication on reasonable request.

## Acknowledgements

SM, AA, and MD thank Dr. Poornima Padmanabhan, Dr. George Thurston, and Dr. Julie Biteen for helpful discussions, and Dr. Mike Norton for detailed discussions at the start of the project. SM also thanks Dr. Omar Saleh for informative discussions. This work was funded by the National Science Foundation Awards DMS-2031179 and DMS-2031180, and by the National Institutes of Health Award 1R01GM143182-01.

