## Supplementary Information for "Morphology, Slow Dynamics, and Composition-Controlled Phase Behavior in a Minimal Model of Dps:DNA Co-Condensation"

#### 1 Ternary Flory–Huggins Free Energy

The Flory–Huggins (FH) theory provides the mean-field framework for describing the thermodynamics of polymer solutions and mixtures. We use it here for an incompressible ternary Dps–DNA–solvent system, with volume fractions $\phi_{\mathrm{Dps}}$, $\phi_{\mathrm{DNA}}$, and $\phi_{\mathrm{Solv}} = 1 - \phi_{\mathrm{Dps}} - \phi_{\mathrm{DNA}}$. In this section we keep the notation general and use the component labels $1, 2, 3$ for DNA, Dps, and solvent, respectively.

The $\phi_{\mathrm{DNA}}$–$\phi_{\mathrm{Dps}}$–$T$ representation of phase behavior provides a useful way to visualize temperature dependence, revealing stability boundaries (spinodals) and coexistence regions (binodals) as surfaces in composition–temperature space. Here we report the formulation, stability criteria, and interpretation of the resulting phase diagrams.

The dimensionless FH free energy density per lattice site can be written as

$$\frac{f}{k_B T} = \frac{\phi_1}{N_1} \ln \phi_1 + \frac{\phi_2}{N_2} \ln \phi_2 + \frac{\phi_3}{N_3} \ln \phi_3 + \chi_{12}\phi_1\phi_2 + \chi_{13}\phi_1\phi_3 + \chi_{23}\phi_2\phi_3, \tag{1}$$

where $N_i$ is the degree of polymerization of component $i$, and $\chi_{ij}$ are the Flory–Huggins interaction parameters.

Each $\chi_{ij}$ is temperature dependent, often expressed as

$$\chi_{ij}(T) = \frac{z}{k_B T} \left( \varepsilon'_{ij} - \tfrac{1}{2}(\varepsilon'_{ii} + \varepsilon'_{jj}) \right), \tag{2}$$

with $z$ the coordination number and $\varepsilon'_{ij}$ the effective pairwise contact energy. Throughout the Flory–Huggins treatment we write these coarse-grained lattice contact energies with a prime ($\varepsilon'$) to distinguish them from the Lennard–Jones well depths $\varepsilon$ used in the Brownian-dynamics simulations (Sec. 9).

#### 2 Thermodynamic Stability: Spinodal Surface

The condition for local stability is that the Hessian $\mathbf{H}$ of the free energy with respect to the independent volume fractions is positive definite. For the ternary case, using $(\phi_1, \phi_2)$ as independent variables (since $\phi_3 = 1 - \phi_1 - \phi_2$), we construct

$$\mathbf{H} = \begin{pmatrix} \frac{\partial^2 f}{\partial \phi_1^2} & \frac{\partial^2 f}{\partial \phi_1 \partial \phi_2} \\ \frac{\partial^2 f}{\partial \phi_2 \partial \phi_1} & \frac{\partial^2 f}{\partial \phi_2^2} \end{pmatrix}. \tag{3}$$

The spinodal surface is defined by the locus of points where

$$\det \mathbf{H} = 0. \tag{4}$$

Inside this surface, the mixture is unstable to infinitesimal fluctuations, whereas outside, the mixture is at least metastable.

### 3 Binodal Surface and Tie–Lines

Phase coexistence (binodal) is determined by the common–tangent construction in free energy space. For two coexisting phases $\alpha$ and $\beta$, the conditions are:

$$\mu_1^{(\alpha)} = \mu_1^{(\beta)}, \tag{5}$$

$$\mu_2^{(\alpha)} = \mu_2^{(\beta)}, \tag{6}$$

$$\Pi^{(\alpha)} = \Pi^{(\beta)}, \tag{7}$$

where $\mu_i = \partial f/\partial \phi_i$ are reduced exchange chemical potentials and $\Pi = \sum_{i=1}^{2} \mu_i \phi_i - f$ is the osmotic pressure in the reduced two-field representation.

These equalities define pairs of compositions $(\phi_1^\alpha, \phi_2^\alpha)$ and $(\phi_1^\beta, \phi_2^\beta)$ at the same temperature $T$, giving the binodal surface. The tie–lines connect coexisting points across this surface.

### 4 Spinodal and Binodal in a Ternary Flory–Huggins Mixture

We consider an incompressible ternary mixture with volume fractions $\phi_1,\ \phi_2,\ \phi_3 = 1 - \phi_1 - \phi_2$, degrees of polymerization $N_1, N_2, N_3$, and temperature–dependent FH parameters $\chi_{ij}(T)$. The free energy density per lattice site (in units of $k_B T$) is

$$\frac{f}{k_B T} = \frac{\phi_1}{N_1}\ln\phi_1 + \frac{\phi_2}{N_2}\ln\phi_2 + \frac{\phi_3}{N_3}\ln\phi_3 + \chi_{12}\phi_1\phi_2 + \chi_{13}\phi_1\phi_3 + \chi_{23}\phi_2\phi_3, \qquad \phi_3 = 1-\phi_1-\phi_2. \tag{8}$$

#### 4.1 Chemical potentials and the Hessian

Because $\phi_3$ is dependent, the two independent variables are $\phi_1, \phi_2$. Define the *reduced* chemical potentials $\mu_1 = \partial f/\partial\phi_1$ and $\mu_2 = \partial f/\partial\phi_2$ at fixed $\phi_2, \phi_1$, respectively (i.e. with $\phi_3 = 1-\phi_1-\phi_2$).

From (8),

$$\mu_1 = k_B T\left[\frac{1}{N_1}(\ln\phi_1 + 1) - \frac{1}{N_3}(\ln\phi_3 + 1) + \chi_{12}\phi_2 + \chi_{13}(\phi_3 - \phi_1) - \chi_{23}\phi_2\right], \tag{9}$$

$$\mu_2 = k_B T\left[\frac{1}{N_2}(\ln\phi_2 + 1) - \frac{1}{N_3}(\ln\phi_3 + 1) + \chi_{12}\phi_1 - \chi_{13}\phi_1 + \chi_{23}(\phi_3 - \phi_2)\right]. \tag{10}$$

Local stability of a homogeneous state requires the $2\times 2$ Hessian $\mathbf{H}$ of second derivatives to be positive definite:

$$\mathbf{H} = \begin{pmatrix} \dfrac{\partial^2 f}{\partial\phi_1^2} & \dfrac{\partial^2 f}{\partial\phi_1\partial\phi_2} \\ \dfrac{\partial^2 f}{\partial\phi_2\partial\phi_1} & \dfrac{\partial^2 f}{\partial\phi_2^2} \end{pmatrix} = k_B T \begin{pmatrix} \dfrac{1}{N_1\phi_1} + \dfrac{1}{N_3\phi_3} - 2\chi_{13} & \dfrac{1}{N_3\phi_3} + \chi_{12} - \chi_{13} - \chi_{23} \\ \dfrac{1}{N_3\phi_3} + \chi_{12} - \chi_{13} - \chi_{23} & \dfrac{1}{N_2\phi_2} + \dfrac{1}{N_3\phi_3} - 2\chi_{23} \end{pmatrix}.$$

#### 4.2 Spinodal surface: $\det\mathbf{H} = 0$

The *spinodal* is the locus where the smallest eigenvalue of $\mathbf{H}$ vanishes, equivalently

$$\det\mathbf{H} = 0 \qquad \Longleftrightarrow \qquad \lambda_{\min}(\mathbf{H}) = 0. \tag{11}$$

For a fixed composition $(\phi_1, \phi_2)$, Eq. (11) is a single equation for the spinodal temperature $T_{\rm sp}(\phi_1, \phi_2)$ because $\chi_{ij} = \chi_{ij}(T)$. For an *UCST* mapping $\chi_{ij}(T) = A_{ij}/T + B_{ij}$, there is typically a largest solution $T_{\rm sp}$ (the one of physical interest); below it $(T < T_{\rm sp})$ the homogeneous state is unstable.

**Critical points on the spinodal.** A critical point (or line) on the spinodal further satisfies vanishing third–order curvature along the unstable eigenvector $\mathbf{v}$ of $\mathbf{H}$:

$$\det \mathbf{H} = 0, \qquad \mathbf{v}^\top \mathbf{H}\, \mathbf{v} = 0, \qquad \mathbf{v}^\top \mathbf{T}[\mathbf{v}, \mathbf{v}] = 0, \tag{12}$$

where $\mathbf{T}$ is the third derivative tensor of $f$ with respect to $(\phi_1, \phi_2)$. In practice, one scans $\det \mathbf{H} = 0$ and tests the condition of unstable eigenvector numerically.

##### 4.3 Binodal (coexistence) surface: common tangent plane

Two phases $\alpha$ and $\beta$ at the same temperature $T$ and compositions $(\phi_1^\alpha, \phi_2^\alpha)$ and $(\phi_1^\beta, \phi_2^\beta)$ coexist when a *single plane* is tangent to $f(\phi_1, \phi_2)$ at both points. Writing the plane as $\ell(\phi_1, \phi_2) = c + m_1\phi_1 + m_2\phi_2$, the tangent conditions are

$$f(\phi_1^\alpha, \phi_2^\alpha) = \ell(\phi_1^\alpha, \phi_2^\alpha), \qquad \nabla f(\phi_1^\alpha, \phi_2^\alpha) = \nabla \ell = (m_1, m_2), \tag{13}$$

$$f(\phi_1^\beta, \phi_2^\beta) = \ell(\phi_1^\beta, \phi_2^\beta), \qquad \nabla f(\phi_1^\beta, \phi_2^\beta) = \nabla \ell = (m_1, m_2). \tag{14}$$

Eliminating $c, m_1, m_2$, the coexistence conditions are the familiar equalities of *two* reduced chemical potentials and the *plane intercept*:

$$\mu_1^{(\alpha)}(\phi^\alpha, T) = \mu_1^{(\beta)}(\phi^\beta, T), \tag{15}$$

$$\mu_2^{(\alpha)}(\phi^\alpha, T) = \mu_2^{(\beta)}(\phi^\beta, T), \tag{16}$$

$$f(\phi^\beta, T) - f(\phi^\alpha, T) = \mu_1^{(\alpha)}(\phi^\alpha, T)\,(\phi_1^\beta - \phi_1^\alpha) + \mu_2^{(\alpha)}(\phi^\alpha, T)\,(\phi_2^\beta - \phi_2^\alpha). \tag{17}$$

Equations (15)–(17) are completely equivalent to equality of *all three* full chemical potentials (including that of component 3) and the osmotic pressure, once incompressibility is accounted for. An alternative but equivalent set uses $\Pi = \sum_{i=1}^{2} \mu_i\phi_i - f$ and imposes $\mu_1^\alpha = \mu_1^\beta$, $\mu_2^\alpha = \mu_2^\beta$, $\Pi^\alpha = \Pi^\beta$.

**Unknowns and solution strategy.** At fixed $T$, (15)–(17) provide three equations for four unknowns $(\phi_1^\alpha, \phi_2^\alpha, \phi_1^\beta, \phi_2^\beta)$. One introduces a one–parameter search (e.g. along an isopleth) or solves the full $(T, \phi^\alpha, \phi^\beta)$ system by continuation. Numerically, a robust approach is:

1. Choose $T$.
2. Minimize the scalar residual $R(\phi^\alpha, \phi^\beta; T) = \|\mu^{(\alpha)} - \mu^{(\beta)}\|_2^2 + \left[f(\phi^\beta) - f(\phi^\alpha) - \mu^{(\alpha)} \cdot (\phi^\beta - \phi^\alpha)\right]^2$ over admissible $\phi^\alpha, \phi^\beta$ (inside the simplex).
3. Discard spurious solutions by requiring both endpoints to lie *outside* the spinodal (i.e. smallest eigenvalue of $\mathbf{H}$ positive).

##### 4.4 Tie–lines and lever rule

For each coexistence pair $(\phi^\alpha, \phi^\beta)$ at temperature $T$, the *tie–line* is the straight segment joining them in the $(\phi_1, \phi_2)$ plane at that $T$. A global composition $\bar{\phi} = (\bar{\phi}_1, \bar{\phi}_2)$ inside the binodal decomposes as

$$\bar{\phi} = \xi\, \phi^\alpha + (1 - \xi)\, \phi^\beta, \qquad 0 \le \xi \le 1, \tag{18}$$

with $\xi$ the phase fraction of the $\alpha$ phase (lever rule). The tie–line direction is the vector $\phi^\beta - \phi^\alpha$ connecting the two coexisting compositions. Near a critical point, this direction approaches the unstable eigenvector associated with the zero mode of the Hessian.

##### 4.5 Summary of implementable conditions

- **Spinodal surface** in $(\phi_1, \phi_2, T)$: solve $\det \mathbf{H}(\phi_1, \phi_2, T) = 0$ for $T$ at each composition; with UCST maps $\chi_{ij}(T) = A_{ij}/T + B_{ij}$, choose the *largest* root.
- **Binodal surface** at fixed $T$: solve (15)–(17) for pairs $(\phi^\alpha, \phi^\beta)$ and retain only endpoints with $\lambda_{\min}(\mathbf{H}) > 0$.
- **Critical set**: points on the spinodal where the third–order directional curvature also vanishes along the null eigenvector.

All required derivatives are explicit in (9)–(10) and in the Hessian above, so no numerical differentiation is needed.

### 5 Mapping of Flory–Huggins Parameters to Pairwise Interaction Energies

In the ternary Flory–Huggins formulation, the temperature dependence of the free energy enters exclusively through the interaction parameters $\chi_{ij}(T)$. These parameters are defined in terms of the underlying lattice contact energies as

$$\chi_{ij}(T) \;=\; \frac{z}{k_B T}\left[\varepsilon'_{ij} - \tfrac{1}{2}\big(\varepsilon'_{ii} + \varepsilon'_{jj}\big)\right], \tag{19}$$

where

- $z$ is the coordination number of the lattice,
- $\varepsilon'_{ij}$ is the nearest–neighbor interaction energy between species $i$ and $j$,
- $\varepsilon'_{ii}, \varepsilon'_{jj}$ are the corresponding like-pair contact energies,
- $k_B T$ is the thermal energy.

The $\varepsilon'_{ij}$ used in this Flory–Huggins mapping are effective lattice contact energies, not the Lennard–Jones well depths used in the Brownian-dynamics simulations. The standard Flory–Huggins parameters are unlike-pair excess interaction parameters. Thus, there are three independent $\chi_{ij}$ values for a ternary mixture: DNA–Dps, DNA–solvent, and Dps–solvent. Like-pair contact energies such as $\varepsilon'_{\mathrm{Dps:Dps}}$ do not give rise to a separate $\chi_{\mathrm{Dps:Dps}}$ parameter; instead, they enter the unlike $\chi_{ij}$ values through the reference term $\frac{1}{2}(\varepsilon'_{ii} + \varepsilon'_{jj})$. Consequently, parameter sweeps shown below in terms of $\varepsilon'_{\mathrm{Dps:DNA}}$ and $\varepsilon'_{\mathrm{Dps:Dps}}$ should be read as sweeps over the underlying contact-energy inputs used to compute the corresponding Flory–Huggins interaction parameters through Eq. (2).

For a ternary mixture of components 1, 2, and 3, this gives three independent interaction parameters:

$$\chi_{12}(T) = \frac{z}{k_B T}\left(\varepsilon'_{12} - \tfrac{1}{2}(\varepsilon'_{11} + \varepsilon'_{22})\right), \tag{20}$$

$$\chi_{13}(T) = \frac{z}{k_B T}\left(\varepsilon'_{13} - \tfrac{1}{2}(\varepsilon'_{11} + \varepsilon'_{33})\right), \tag{21}$$

$$\chi_{23}(T) = \frac{z}{k_B T}\left(\varepsilon'_{23} - \tfrac{1}{2}(\varepsilon'_{22} + \varepsilon'_{33})\right). \tag{22}$$

These definitions quantify the excess energetic cost of unlike contacts relative to the mean of like–like interactions:

- If $\varepsilon'_{ij} \approx \frac{1}{2}(\varepsilon'_{ii} + \varepsilon'_{jj})$, then $\chi_{ij} \approx 0$, corresponding to nearly ideal mixing.
- If $\varepsilon'_{ij}$ is higher than the mean of the like–like contacts, then $\chi_{ij} > 0$, favoring demixing.
- If $\varepsilon'_{ij}$ is lower, then $\chi_{ij} < 0$, favoring attractive mixing.

The ternary Flory–Huggins free energy density (per lattice site, in units of $k_B T$) can then be written as

$$\frac{f}{k_B T} = \frac{\phi_1}{N_1} \ln \phi_1 + \frac{\phi_2}{N_2} \ln \phi_2 + \frac{\phi_3}{N_3} \ln \phi_3 + \chi_{12}(T)\phi_1\phi_2 + \chi_{13}(T)\phi_1\phi_3 + \chi_{23}(T)\phi_2\phi_3, \tag{23}$$

where $\phi_i$ are the volume fractions and $N_i$ the degrees of polymerization of components $i = 1, 2, 3$.

### 6 Temperature sensitivity via binary Flory–Huggins slices

The full ternary FH free energy (8) depends on two independent composition variables $(\phi_1, \phi_2)$ and on temperature through the interaction parameters $\chi_{ij}(T)$. As a result, both the spinodal and the binodal are three–dimensional surfaces in $(\phi_1, \phi_2, T)$ space. While this representation is complete, it can be difficult to visualize the effect of temperature alone. To highlight the sensitivity of phase behavior to $T$ in a simpler way, it is convenient to consider *effective binary cuts* through the ternary diagram and construct the associated composition–temperature phase diagrams.

#### 6.1 Effective binary free energy

Along a given path in the ternary diagram (for example, at fixed solvent content $\phi_3$ or along a composition line that primarily varies one of the non-solvent components), the ternary free energy can be reduced to an effective binary FH form. Writing $\phi$ for the volume fraction of one effective component along such a path, we obtain schematically

$$\frac{f_{\text{bin}}}{k_B T} = \frac{\phi}{N_1} \ln \phi + \frac{1-\phi}{N_2} \ln(1-\phi) + \chi(T)\,\phi(1-\phi), \tag{24}$$

where $N_1$ and $N_2$ are effective degrees of polymerization for the two species and $\chi(T)$ is an effective interaction parameter that inherits the $1/T$ dependence from the underlying $\chi_{ij}(T)$,

$$\chi(T) = \frac{\chi_0}{T}, \tag{25}$$

with $\chi_0$ determined by the microscopic contact energies $\varepsilon'_{ij}$ entering the ternary model. This binary representation is not meant to describe any single cut of the ternary diagram in detail, but rather to capture the generic way in which temperature controls the competition between entropic and enthalpic contributions.

#### 6.2 Spinodal, binodal, and critical point in the $\phi$–$T$ plane

For the binary free energy (24), the spinodal is defined by the loss of convexity,

$$\frac{\partial^2 f_{\text{bin}}}{\partial \phi^2} = 0, \tag{26}$$

which yields the well-known FH expression

$$\chi_{\text{sp}}(\phi) = \frac{1}{2}\left(\frac{1}{N_1\phi} + \frac{1}{N_2(1-\phi)}\right). \tag{27}$$

Using $\chi(T) = \chi_0/T$, this relation defines a spinodal curve $T_{\mathrm{sp}}(\phi)$ in the $\phi$–$T$ plane.

The binodal (coexistence) curve consists of pairs of compositions $(\phi_1, \phi_2)$ at a given temperature $T$ that satisfy the common–tangent conditions

$$\mu(\phi_1, T) = \mu(\phi_2, T), \tag{28}$$

$$f_{\mathrm{bin}}(\phi_2, T) - f_{\mathrm{bin}}(\phi_1, T) = \mu(\phi_1, T)\,(\phi_2 - \phi_1), \tag{29}$$

where $\mu(\phi, T) = \partial f_{\mathrm{bin}}/\partial\phi$ is the reduced chemical potential. These conditions are the binary analogue of Eqs. (15)–(17) and define the red binodal dome in the $\phi$–$T$ diagrams shown in Fig. 1. For each temperature $T < T_c$, the tie–line connects the two coexisting compositions $\phi_1(T)$ and $\phi_2(T)$ at that temperature.

The critical point $(\phi_c, T_c)$ for the binary FH model can be obtained analytically,

$$\phi_c = \frac{\sqrt{N_2}}{\sqrt{N_1} + \sqrt{N_2}}, \qquad \chi_c = \frac{1}{2}\left(\frac{1}{\sqrt{N_1}} + \frac{1}{\sqrt{N_2}}\right)^2, \qquad T_c = \frac{\chi_0}{\chi_c}. \tag{30}$$

For a symmetric mixture $N_1 = N_2$, one recovers the symmetric critical composition $\phi_c = 1/2$ and a binodal that is symmetric about $\phi = 1/2$. When $N_2 \neq N_1$, both the critical composition and the shape of the binodal become asymmetric, as discussed below.

#### 6.3 Symmetric versus asymmetric mixtures

Figure 1 shows three examples of the binary $\phi$–$T$ diagram obtained from the effective FH model (24): a symmetric mixture with $N_1 = N_2 = 1$ and two increasingly asymmetric mixtures with $N_1 = 1$, $N_2 = 3$ and $N_1 = 1$, $N_2 = 10$. In each panel, the blue dots denote the spinodal, the red dots trace the binodal dome, gray lines indicate tie–lines at selected temperatures, and the black dot marks the critical point.

In the symmetric case ($N_1 = N_2$), the binodal and spinodal are symmetric around $\phi_c = 1/2$. As temperature decreases from high values, the homogeneous mixture first becomes metastable (crossing the binodal) and eventually unstable (crossing the spinodal). The tie–lines span a wide range of compositions, illustrating how a given overall composition inside the dome decomposes into two coexisting phases, one enriched in component 1 and the other enriched in component 2.

As the mixture becomes asymmetric ($N_2 > N_1$), the critical value of the plotted composition $\phi$ shifts to larger values; equivalently, the critical volume fraction of the longer component, $1-\phi_c$, decreases. The binodal becomes skewed and the coexistence region in the $\phi$–$T$ plane narrows on one side. Consequently, the range of temperatures and compositions that admit two distinct coexisting phases becomes more restricted, and the tie–lines shrink or disappear except near the critical point. The critical temperature shifts to higher values for longer polymers. Because the translational entropy of the polymeric component is weakened by the $1/N$ scaling in Eq. 1, phase separation can occur at smaller effective interaction parameters $\chi$. For $\chi(T) \propto 1/T$, this corresponds to a higher critical temperature. This behavior is a direct consequence of the FH expressions for $\phi_c$ and $\chi_c$ and is a generic feature of polymer–solvent or polymer–polymer mixtures with asymmetric compositions.

#### 6.4 Relevance for the ternary Dps–DNA–solvent model

Although Fig. 1 is based on an effective binary representation, it captures the essential way in which temperature controls phase separation in the full ternary Dps–DNA–solvent system. In the ternary case, each point in the $\phi_1$–$\phi_2$ plane can be associated with an effective binary cut of the type described above, with an interaction parameter $\chi(T)$ inherited from the underlying $\chi_{ij}(T)$. The binary domes in Fig. 1 can therefore be viewed as schematic slices through the full $\phi_1$–$\phi_2$–$T$ phase diagram discussed in the previous sections.

In the main text, we focus on a fixed effective temperature (i.e. a fixed set of $\chi_{ij}$) to study how Dps–DNA condensates form and reorganize as the composition and interaction strengths are varied. The binary $\phi$–$T$ diagrams here serve to illustrate that, within the same FH framework, lowering $T$ (or equivalently increasing the effective $\chi_{ij}$) would generically enlarge the two-phase region, shift the critical composition, and modify the coexistence window. Thus, even though temperature is held fixed in our simulation study, the underlying mean-field theory predicts a generic sensitivity of this phase behavior to $T$.

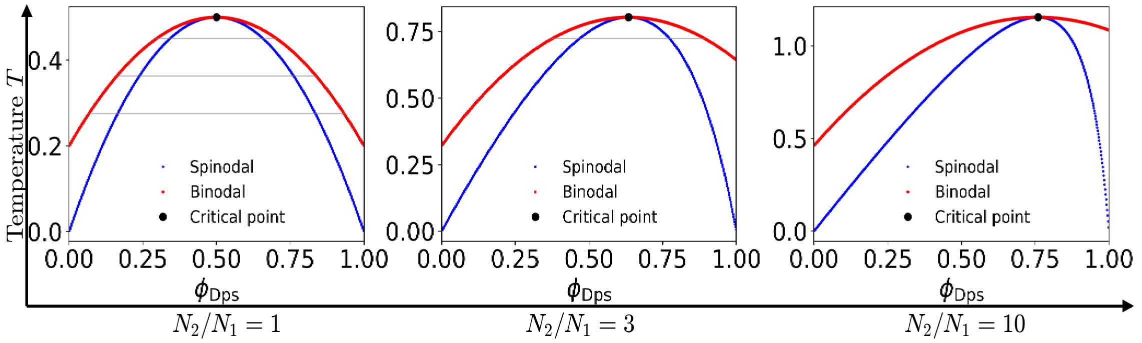


Figure 1: Binary Flory–Huggins phase diagrams illustrating temperature dependence. Each panel shows the spinodal (blue), binodal (red), tie–lines (gray), and critical point (black) in the composition–temperature plane for an effective binary mixture, with the critical point given by Eq. 30. (Left) Symmetric mixture $N_1 = N_2 = 1$ with critical composition $\phi_c = 1/2$. (Middle, right) Asymmetric mixtures with $N_1 = 1$, $N_2 = 3$ and $N_1 = 1$, $N_2 = 10$, respectively, for which the critical composition shifts and the coexistence window becomes increasingly skewed and narrow. These diagrams serve as schematic binary slices through the full ternary Dps–DNA–solvent phase landscape, highlighting the generic sensitivity of phase separation to temperature.

#### 7 Ternary phase diagrams

An incompressible ternary mixture is completely described by two independent volume fractions, since the third is fixed by the incompressibility constraint. In the Dps–DNA–solvent system, each composition may therefore be represented either on the reduced $(\phi_{\text{DNA}}, \phi_{\text{Dps}})$ plane, with $\phi_{\text{Solv}} = 1 - \phi_{\text{Dps}} - \phi_{\text{DNA}}$, or equivalently on a ternary diagram whose three axes are $\phi_{\text{Dps}}$, $\phi_{\text{DNA}}$, and $\phi_{\text{Solv}}$.

To examine how the coexistence region changes with interaction strength, we varied the underlying pairwise contact-energy inputs and recomputed the Flory–Huggins phase diagram at each parameter set. Figures 2 and 3 show the same sweep in two complementary views: Fig. 2 uses the reduced binary-style projection, while Fig. 3 redraws the same phase behavior on ternary composition triangles to make the solvent coordinate explicit. The axes are shown in terms of $\varepsilon'$ because the varied quantities are contact-energy inputs to the Flory–Huggins mapping, not independent $\chi$ parameters; in particular, $\varepsilon'_{\text{Dps:Dps}}$ enters the effective unlike interaction parameters, such as $\chi_{\text{Dps:DNA}}$ and $\chi_{\text{Dps:Solv}}$, through the reference-energy terms in Eq. (2).

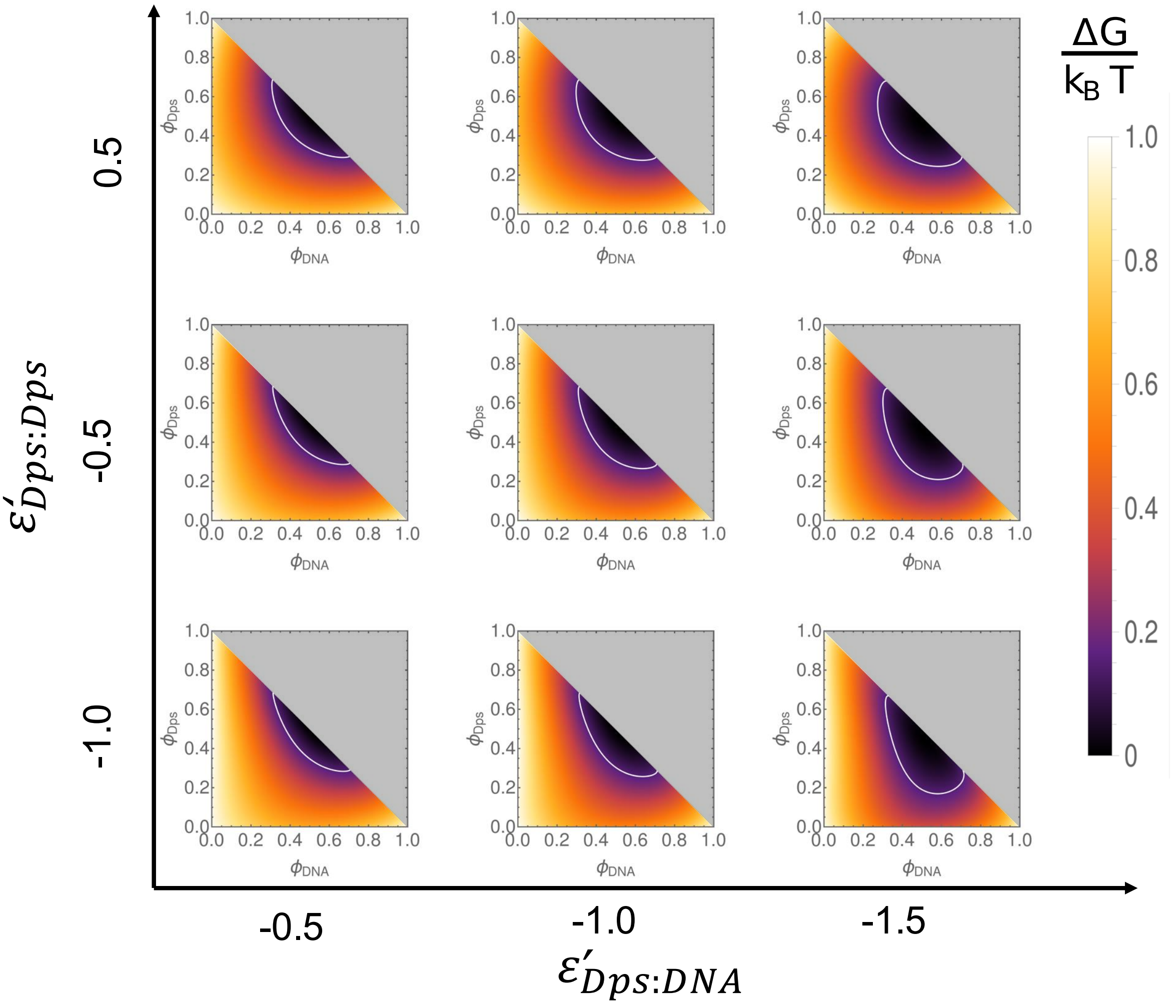


Figure 2: Reduced binary-style representation of the contact-energy sweep for the ternary Dps–DNA–solvent Flory–Huggins model. Each panel shows the incompressible composition plane using the two independent coordinates $(\phi_{\mathrm{DNA}}, \phi_{\mathrm{Dps}})$, with the solvent fraction fixed by $\phi_{\mathrm{Solv}} = 1 - \phi_{\mathrm{Dps}} - \phi_{\mathrm{DNA}}$. The panels correspond to the indicated values of $\varepsilon'_{\mathrm{Dps:DNA}}$ and $\varepsilon'_{\mathrm{Dps:Dps}}$, and the corresponding Flory–Huggins parameters $\chi_{ij}$ are obtained from Eq. (2). The white contour denotes the binodal; the enclosed region identifies the coexistence regime where a Dps–DNA-rich condensed phase coexists with a solvent-rich phase. The gray triangular regions are compositionally inaccessible under the incompressibility constraint. The plotted free energy density is rescaled to lie bewteen 0 and 1.

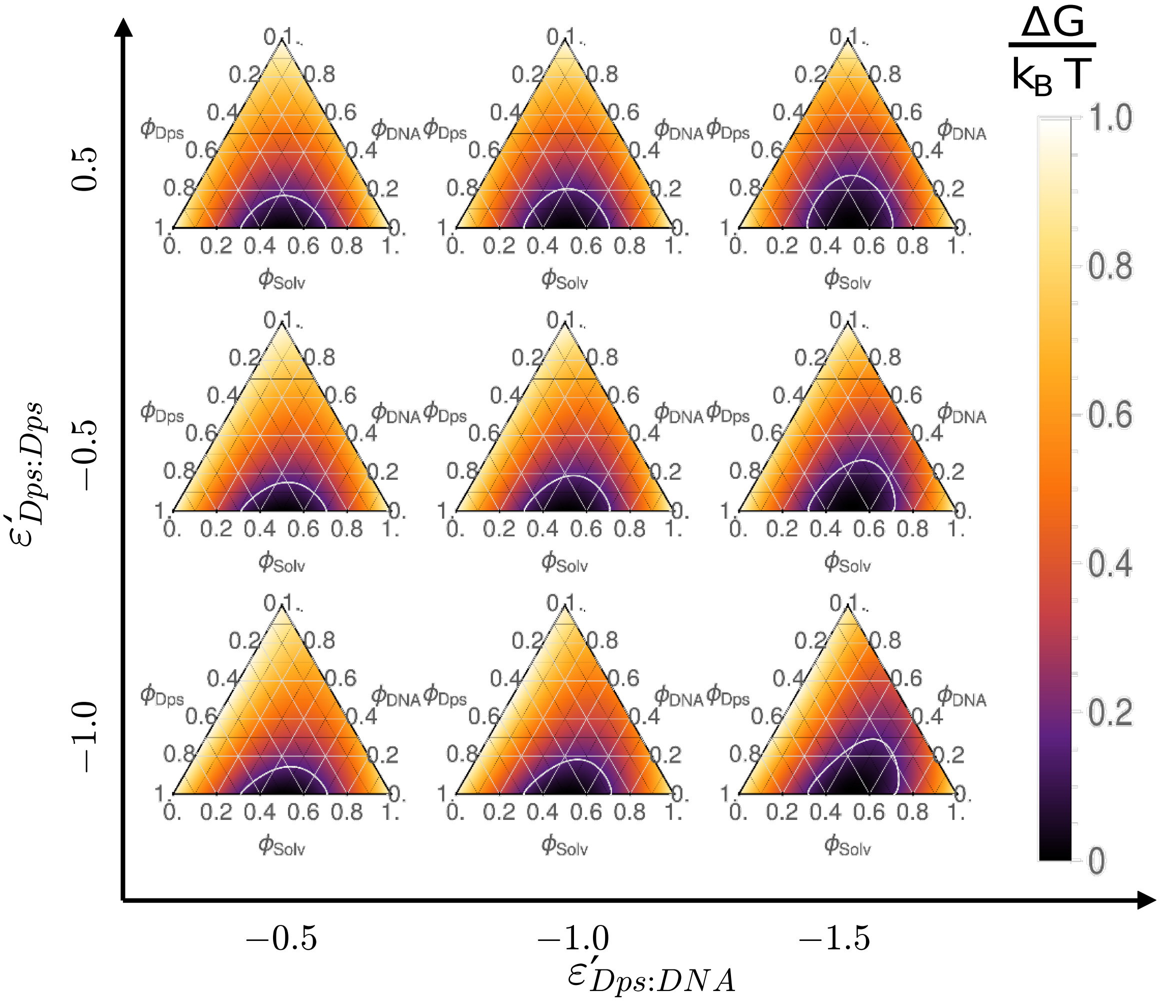


Figure 3: Ternary-coordinate representation of the same contact-energy sweep shown in Fig. 2. Each small triangle shows the Dps–DNA–solvent composition plane for the same values of $\varepsilon'_{\mathrm{Dps:DNA}}$ and $\varepsilon'_{\mathrm{Dps:Dps}}$, with the solvent coordinate shown explicitly through the ternary axes. Pairwise contact energies involving solvent were fixed at $\varepsilon'_{\mathrm{DNA:Solv}} = \varepsilon'_{\mathrm{Dps:Solv}} = \varepsilon'_{\mathrm{Solv:Solv}} = -0.1$, and the DNA–DNA contact energy was held fixed. The white contour denotes the same binodal construction as in Fig. 2; the enclosed region identifies the coexistence regime where a Dps–DNA-rich condensed phase coexists with a solvent-rich phase. The plotted free energy density is rescaled to lie bewteen 0 and 1.

### 8 Ternary Flory–Huggins–Cahn–Hilliard–Cook simulations

#### 8.1 Composition variables and incompressibility

We model the Dps:DNA:solvent mixture as an incompressible ternary composition field. The local volume fractions of Dps, DNA, and solvent are denoted by

$$\phi_{\mathrm{Dps}}(\mathbf{r},t), \qquad \phi_{\mathrm{DNA}}(\mathbf{r},t), \qquad \phi_{\mathrm{Solv}}(\mathbf{r},t),$$

with the incompressibility constraint

$$\phi_{\mathrm{Dps}} + \phi_{\mathrm{DNA}} + \phi_{\mathrm{Solv}} = 1. \tag{31}$$

Thus only two fields are independent. We evolve $\phi_{\mathrm{Dps}}$ and $\phi_{\mathrm{DNA}}$ explicitly, and compute the solvent field from

$$\phi_{\mathrm{Solv}} = 1 - \phi_{\mathrm{Dps}} - \phi_{\mathrm{DNA}}. \tag{32}$$

The total solute or co-condensate field is

$$\phi_T = \phi_{\mathrm{Dps}} + \phi_{\mathrm{DNA}}. \tag{33}$$

This field is useful for identifying Dps:DNA-rich and solvent-rich regions, although the simulations retain the two independent solute fields separately.

#### 8.2 Local Flory–Huggins free-energy density

The local mixing free-energy density is written in dimensionless form, in units of $k_BT$, as

$$\begin{aligned} f_{\mathrm{FH}} = {} & \frac{\phi_{\mathrm{DNA}}}{N_{\mathrm{DNA}}}\ln\phi_{\mathrm{DNA}} + \frac{\phi_{\mathrm{Dps}}}{N_{\mathrm{Dps}}}\ln\phi_{\mathrm{Dps}} + \frac{\phi_{\mathrm{Solv}}}{N_{\mathrm{Solv}}}\ln\phi_{\mathrm{Solv}} \\ & + \chi_{\mathrm{Dps:DNA}}\phi_{\mathrm{Dps}}\phi_{\mathrm{DNA}} + \chi_{\mathrm{Dps:Solv}}\phi_{\mathrm{Dps}}\phi_{\mathrm{Solv}} + \chi_{\mathrm{DNA:Solv}}\phi_{\mathrm{DNA}}\phi_{\mathrm{Solv}}. \end{aligned} \tag{34}$$

Here $N_{\mathrm{Dps}}$, $N_{\mathrm{DNA}}$, and $N_{\mathrm{Solv}}$ are the effective polymerization indices of the three components. In the phase-field calculations reported for the morphology plots, we used

$$N_{\mathrm{Dps}} = 1, \qquad N_{\mathrm{Solv}} = 1, \qquad N_{\mathrm{DNA}} = 100.$$

Thus Dps and solvent are treated as small-molecule components, whereas DNA has reduced translational entropy through the polymeric $\phi_{\mathrm{DNA}}\ln\phi_{\mathrm{DNA}}/N_{\mathrm{DNA}}$ term. The value of $N_{\mathrm{DNA}}$ should be interpreted as an effective coarse-grained polymerization parameter for the continuum model, rather than as the number of beads in the Brownian dynamics simulations.

Using the incompressibility constraint, the solvent field is eliminated and the free-energy density is reduced to a function of the two independent fields:

$$f(\phi_{\mathrm{DNA}}, \phi_{\mathrm{Dps}}) = f_{\mathrm{FH}}\left(\phi_{\mathrm{DNA}}, \phi_{\mathrm{Dps}}, 1 - \phi_{\mathrm{DNA}} - \phi_{\mathrm{Dps}}\right). \tag{35}$$

#### 8.3 Exchange chemical potentials

Because the solvent is eliminated by incompressibility, the relevant driving forces are exchange chemical potentials relative to the solvent. The local parts of these chemical potentials are

$$g_i = \frac{\partial f}{\partial \phi_i}, \qquad i \in \{\mathrm{Dps}, \mathrm{DNA}\}, \tag{36}$$

where the derivative is taken at fixed value of the other independent field.

Writing $\phi_{\text{Solv}} = 1 - \phi_{\text{Dps}} - \phi_{\text{DNA}}$, the explicit local exchange chemical potentials are

$$g_{\text{Dps}} = \frac{\ln \phi_{\text{Dps}} + 1}{N_{\text{Dps}}} - \frac{\ln \phi_{\text{Solv}} + 1}{N_{\text{Solv}}} + \chi_{\text{Dps:DNA}}\phi_{\text{DNA}} + \chi_{\text{Dps:Solv}}\left(\phi_{\text{Solv}} - \phi_{\text{Dps}}\right) - \chi_{\text{DNA:Solv}}\phi_{\text{DNA}}, \tag{37}$$

$$g_{\text{DNA}} = \frac{\ln \phi_{\text{DNA}} + 1}{N_{\text{DNA}}} - \frac{\ln \phi_{\text{Solv}} + 1}{N_{\text{Solv}}} + \chi_{\text{Dps:DNA}}\phi_{\text{Dps}} - \chi_{\text{Dps:Solv}}\phi_{\text{Dps}} + \chi_{\text{DNA:Solv}}\left(\phi_{\text{Solv}} - \phi_{\text{DNA}}\right). \tag{38}$$

In the numerical implementation, the logarithms are regularized by replacing each volume fraction inside the logarithm by $\max(\phi_i, \varepsilon_{\log})$, with

$$\varepsilon_{\log} = 10^{-7}.$$

#### 8.4 Square-gradient free energy

To describe finite-width interfaces between Dps:DNA-rich and solvent-rich regions, we add square-gradient penalties to the local Flory–Huggins free energy. The dimensionless free-energy functional is

$$\frac{F}{k_B T} = \int d\mathbf{r} \Big[ f(\phi_{\text{DNA}}, \phi_{\text{Dps}}) + \frac{\kappa_{\text{DNA}}}{2}|\nabla\phi_{\text{DNA}}|^2 + \frac{\kappa_{\text{Dps}}}{2}|\nabla\phi_{\text{Dps}}|^2 + \kappa_{\text{Dps:DNA}}\nabla\phi_{\text{Dps}} \cdot \nabla\phi_{\text{DNA}} \Big]. \tag{39}$$

The coefficients $\kappa_{\text{Dps}}$, $\kappa_{\text{DNA}}$, and $\kappa_{\text{Dps:DNA}}$ set the interfacial width and curvature cost. The cross-gradient term allows the two solute fields to share interfaces and co-localize within a common dense phase.

The gradient contribution is stable when the gradient-penalty matrix is positive definite. For the symmetric case $\kappa_{\text{Dps}} = \kappa_{\text{DNA}}$, this requires

$$\kappa^2_{\text{Dps:DNA}} < \kappa_{\text{Dps}}\kappa_{\text{DNA}}. \tag{40}$$

All parameter choices used below satisfy this condition.

The full variational chemical potentials are

$$\mu_{\text{Dps}} = \frac{\delta F}{\delta \phi_{\text{Dps}}} = g_{\text{Dps}} - \kappa_{\text{Dps}}\nabla^2\phi_{\text{Dps}} - \kappa_{\text{Dps:DNA}}\nabla^2\phi_{\text{DNA}}, \tag{41}$$

$$\mu_{\text{DNA}} = \frac{\delta F}{\delta \phi_{\text{DNA}}} = g_{\text{DNA}} - \kappa_{\text{DNA}}\nabla^2\phi_{\text{DNA}} - \kappa_{\text{Dps:DNA}}\nabla^2\phi_{\text{Dps}}. \tag{42}$$

#### 8.5 Conserved Cahn–Hilliard–Cook dynamics

The two independent conserved composition fields evolve according to Cahn–Hilliard–Cook dynamics:

$$\partial_t \phi_i = \nabla \cdot (M_i \nabla \mu_i) + \nabla \cdot \boldsymbol{\eta}_i, \qquad i \in \{\text{Dps}, \text{DNA}\}. \tag{43}$$

Here $M_i$ are the mobilities, $\mu_i$ are the variational chemical potentials in Eqs. (41) and (42), and $\boldsymbol{\eta}_i$ is a conserved stochastic flux noise. The divergence form ensures that the spatial integral of each independent field is conserved. The solvent fraction then follows from Eq. (32).

For an ideal continuum CHC model, the stochastic flux noise satisfies the fluctuation–dissipation structure

$$\left\langle \eta_{i\alpha}(\mathbf{r}, t)\eta_{j\beta}(\mathbf{r}', t')\right\rangle = 2k_B T M_i \delta_{ij}\delta_{\alpha\beta}\delta(\mathbf{r} - \mathbf{r}')\delta(t - t'), \tag{44}$$

where $\alpha$ and $\beta$ denote Cartesian components. In the dimensionless numerical simulations used here, the noise was implemented as a weak conserved flux perturbation, described below, providing weak stochastic seeding together with a small ongoing conserved perturbation during the evolution. The morphology classification does not rely on the quantitative amplitude of thermal fluctuations.

##### 8.6 Numerical domain and spectral discretization

The simulations were performed on a two-dimensional periodic square domain of side length

$$L = 128$$

with

$$N_x = N_y = 128, \qquad \Delta x = \Delta y = L/N_x = 1.$$

The timestep was

$$\Delta t = 0.030.$$

Periodic boundary conditions were used in both spatial directions.

Spatial derivatives were evaluated pseudospectrally. The Fourier wavenumbers were

$$k_x = \frac{2\pi}{L}\left(0, 1, \ldots, \frac{N}{2} - 1, -\frac{N}{2}, \ldots, -1\right), \tag{45}$$

with an analogous expression for $k_y$. Defining

$$k^2 = k_x^2 + k_y^2, \qquad k^4 = (k^2)^2,$$

the Laplacian and biharmonic operators are diagonal in Fourier space.

The local chemical potentials $g_{\mathrm{Dps}}$ and $g_{\mathrm{DNA}}$ were evaluated in real space from Eqs. (37) and (38), transformed to Fourier space, and used in a semi-implicit update. The gradient terms were treated implicitly, while the nonlinear local Flory–Huggins terms were treated explicitly.

For a Fourier mode with wavenumber magnitude $k$, the update has the linear system

$$D_{\mathrm{Dps,Dps}}\,\widehat{\phi}_{\mathrm{Dps}}^{n+1} + D_{\mathrm{Dps,DNA}}\,\widehat{\phi}_{\mathrm{DNA}}^{n+1} = R_{\mathrm{Dps}}, \tag{46}$$

$$D_{\mathrm{DNA,Dps}}\,\widehat{\phi}_{\mathrm{Dps}}^{n+1} + D_{\mathrm{DNA,DNA}}\,\widehat{\phi}_{\mathrm{DNA}}^{n+1} = R_{\mathrm{DNA}}, \tag{47}$$

where

$$D_{\mathrm{Dps,Dps}} = 1 + \Delta t\, M_{\mathrm{Dps}} S_{\mathrm{Dps}} k^2 + \Delta t\, M_{\mathrm{Dps}} \kappa_{\mathrm{Dps}} k^4, \tag{48}$$

$$D_{\mathrm{DNA,DNA}} = 1 + \Delta t\, M_{\mathrm{DNA}} S_{\mathrm{DNA}} k^2 + \Delta t\, M_{\mathrm{DNA}} \kappa_{\mathrm{DNA}} k^4, \tag{49}$$

$$D_{\mathrm{Dps,DNA}} = \Delta t\, M_{\mathrm{Dps}} \kappa_{\mathrm{Dps:DNA}} k^4, \tag{50}$$

$$D_{\mathrm{DNA,Dps}} = \Delta t\, M_{\mathrm{DNA}} \kappa_{\mathrm{Dps:DNA}} k^4. \tag{51}$$

The right-hand sides are

$$R_{\mathrm{Dps}} = \left(1 + \Delta t\, M_{\mathrm{Dps}} S_{\mathrm{Dps}} k^2\right) \widehat{\phi}_{\mathrm{Dps}}^{n} - \Delta t\, M_{\mathrm{Dps}} k^2 \widehat{g}_{\mathrm{Dps}}^{n}, \tag{52}$$

$$R_{\mathrm{DNA}} = \left(1 + \Delta t\, M_{\mathrm{DNA}} S_{\mathrm{DNA}} k^2\right) \widehat{\phi}_{\mathrm{DNA}}^{n} - \Delta t\, M_{\mathrm{DNA}} k^2 \widehat{g}_{\mathrm{DNA}}^{n}. \tag{53}$$

The constants

$$S_{\mathrm{Dps}} = S_{\mathrm{DNA}} = 350$$

are linear stabilization parameters. They improve numerical stability for the explicit local free-energy terms and do not alter the conserved structure of the dynamics.

For each Fourier mode, the $2 \times 2$ system is solved analytically:

$$\widehat{\phi}_{\mathrm{Dps}}^{n+1} = \frac{R_{\mathrm{Dps}} D_{\mathrm{DNA,DNA}} - D_{\mathrm{Dps,DNA}} R_{\mathrm{DNA}}}{D_{\mathrm{Dps,Dps}} D_{\mathrm{DNA,DNA}} - D_{\mathrm{Dps,DNA}} D_{\mathrm{DNA,Dps}}}, \tag{54}$$

$$\widehat{\phi}_{\mathrm{DNA}}^{n+1} = \frac{D_{\mathrm{Dps,Dps}} R_{\mathrm{DNA}} - D_{\mathrm{DNA,Dps}} R_{\mathrm{Dps}}}{D_{\mathrm{Dps,Dps}} D_{\mathrm{DNA,DNA}} - D_{\mathrm{Dps,DNA}} D_{\mathrm{DNA,Dps}}}. \tag{55}$$

The updated fields are then transformed back to real space.

#### 8.7 Weak conserved Cook noise implementation

For the Cahn–Hilliard–Cook simulations, the stochastic term was added in conserved form as the divergence of a random flux. At each timestep and for each independent field, independent Gaussian random fluxes $J_x(\mathbf{r})$ and $J_y(\mathbf{r})$ were generated on the periodic grid. The scalar conserved noise increment was computed as the discrete periodic divergence

$$\xi(\mathbf{r}) = J_x(\mathbf{r}) - J_x(\mathbf{r} - \Delta x\,\hat{\mathbf{x}}) + J_y(\mathbf{r}) - J_y(\mathbf{r} - \Delta y\,\hat{\mathbf{y}}). \tag{56}$$

Because this is a periodic divergence, its spatial sum vanishes up to roundoff:

$$\sum_{\mathbf{r}} \xi(\mathbf{r}) = 0. \tag{57}$$

The noise field was normalized to unit standard deviation and added as

$$\phi_{\mathrm{Dps}}^{n+1} \leftarrow \phi_{\mathrm{Dps}}^{n+1} + \sigma_{\mathrm{C}} \xi_{\mathrm{Dps}}, \tag{58}$$

$$\phi_{\mathrm{DNA}}^{n+1} \leftarrow \phi_{\mathrm{DNA}}^{n+1} + \sigma_{\mathrm{C}} \xi_{\mathrm{DNA}}. \tag{59}$$

The weak Cook-noise amplitude used for morphology calculations was

$$\sigma_{\mathrm{C}} = 10^{-5}. \tag{60}$$

We also checked weaker values, including $\sigma_{\mathrm{C}} = 10^{-6}$, and the qualitative morphology classification was unchanged. The Cook noise therefore serves as a weak conserved stochastic perturbation and does not set the observed droplet/network morphology.

#### 8.8 Initial conditions

Each simulation was initialized near a homogeneous state with prescribed mean compositions

$$\bar{\phi}_{\mathrm{Dps}}, \qquad \bar{\phi}_{\mathrm{DNA}}.$$

Smooth correlated initial perturbations were added to seed phase separation. A common Gaussian random field $\zeta_{\mathrm{com}}$ and a weak anti-correlated Gaussian random field $\zeta_{\mathrm{anti}}$ were generated and smoothed in Fourier space with the filter

$$\widehat{\zeta}(\mathbf{k}) \leftarrow \widehat{\zeta}(\mathbf{k}) \exp\left(-\frac{1}{2}\sigma_{\mathrm{init}}^2 k^2\right). \tag{61}$$

Each smoothed field was normalized to unit standard deviation. The initial fields were then set to

$$\phi_{\mathrm{Dps}}(\mathbf{r}, 0) = \bar{\phi}_{\mathrm{Dps}} + A_{\mathrm{init}} \zeta_{\mathrm{com}} + 0.06 A_{\mathrm{init}} \zeta_{\mathrm{anti}}, \tag{62}$$

$$\phi_{\mathrm{DNA}}(\mathbf{r}, 0) = \bar{\phi}_{\mathrm{DNA}} + A_{\mathrm{init}} \left(\frac{\bar{\phi}_{\mathrm{DNA}}}{\bar{\phi}_{\mathrm{Dps}}}\right) \zeta_{\mathrm{com}} - 0.04 A_{\mathrm{init}} \left(\frac{\bar{\phi}_{\mathrm{DNA}}}{\bar{\phi}_{\mathrm{Dps}}}\right) \zeta_{\mathrm{anti}}. \tag{63}$$

The common component biases Dps and DNA to co-condense into the same dense phase, while the weak anti-correlated component allows local composition variations between the two solutes. The initial noise amplitude was

$$A_{\text{init}} = 4 \times 10^{-3}.$$

For the balanced Dps:DNA case, the smoothing length was

$$\sigma_{\text{init}} = 5,$$

whereas for the Dps-rich/DNA-lower droplet case,

$$\sigma_{\text{init}} = 10.$$

The larger initial smoothing length in the droplet case suppresses grid-scale nuclei and gives a cleaner long-wavelength droplet morphology.

##### 8.9 Conservative projection onto the physical simplex

After initialization and after each timestep, the fields were projected back into the physical ternary simplex. The projection enforces

$$\phi_{\text{Dps}} \geq \varepsilon_{\log}, \qquad \phi_{\text{DNA}} \geq \varepsilon_{\log}, \qquad \phi_{\text{Dps}} + \phi_{\text{DNA}} \leq 1 - \varepsilon_{\log},$$

while preserving the prescribed spatial means $\bar{\phi}_{\text{Dps}}$ and $\bar{\phi}_{\text{DNA}}$.

The projection was implemented iteratively. First, each field was shifted by a spatially uniform amount to restore its target mean. Second, negative or near-zero values were clipped to $\varepsilon_{\log}$. Third, at grid points where $\phi_{\text{Dps}} + \phi_{\text{DNA}} > 1 - \varepsilon_{\log}$, both solute fields were rescaled proportionally so that the total solute fraction satisfied the incompressibility bound. This procedure was repeated 24 times per projection call. This projection removes small roundoff and overshoot errors while retaining conservation of the two independent species.

##### 8.10 Parameter sets used for morphology simulations

The morphology simulations in the phase-field model were specified directly in terms of the effective Flory–Huggins interaction parameters. These values are dimensionless coarse-grained thermodynamic parameters and should not be identified with the Brownian-dynamics Lennard–Jones well depths. The same Flory–Huggins interaction parameters were used in both composition rows:

$$\chi_{\text{Dps:DNA}} = -2.0, \qquad \chi_{\text{Dps:Solv}} = 2.0, \qquad \chi_{\text{DNA:Solv}} = 2.0. \tag{64}$$

The negative $\chi_{\text{Dps:DNA}}$ favors Dps:DNA association, while the positive solute–solvent parameters favor demixing from the solvent.

The droplet-like case used mean composition

$$(\bar{\phi}_{\text{Dps}}, \bar{\phi}_{\text{DNA}}) = (0.30, 0.10), \tag{65}$$

with interfacial penalties

$$\kappa_{\text{Dps}} = 2.4, \qquad \kappa_{\text{DNA}} = 2.4, \qquad \kappa_{\text{Dps:DNA}} = 1.8. \tag{66}$$

This Dps-rich/DNA-lower composition coarsens into isolated Dps:DNA-rich droplets coexisting with a solvent-rich background.

The network-like case used mean composition

$$(\bar{\phi}_{\text{Dps}}, \bar{\phi}_{\text{DNA}}) = (0.21, 0.21), \tag{67}$$

with interfacial penalties

$$\kappa_{\mathrm{Dps}} = 1.1, \qquad \kappa_{\mathrm{DNA}} = 1.1, \qquad \kappa_{\mathrm{Dps:DNA}} = 0.85. \tag{68}$$

This balanced Dps:DNA composition coarsens into an interconnected, bicontinuous Dps:DNA-rich morphology.

For the final equal-mobility calculations, we used

$$M_{\mathrm{Dps}} = M_{\mathrm{DNA}} = 1. \tag{69}$$

We also tested unequal mobilities and found that the morphology classes at the saved times were unchanged. Thus, within the parameter range used here, the mobilities primarily set the kinetic clock rather than selecting between droplet-like and network-like morphologies.

##### 8.11 Role of the interfacial parameters

The square-gradient coefficients are effective dimensionless parameters that set the interfacial width, curvature penalty, and coarsening length scale in the phase-field model. They are not interpreted as directly measured material constants. Smaller $\kappa$ values produce sharper interfaces and finer domains, whereas larger $\kappa$ values suppress short-wavelength structure and produce smoother domains.

The qualitative droplet/network distinction is controlled primarily by the mean composition and Flory–Huggins interaction parameters. The interfacial parameters tune how this thermodynamic instability is resolved on a finite grid. In particular, the Dps-rich/DNA-lower composition favors isolated dense domains, while the balanced Dps:DNA composition favors extended bicontinuous connectivity. Reducing the $\kappa$ values shifts the apparent coarsening times and sharpens interfaces, but does not remove the qualitative morphology distinction.

##### 8.12 Saved fields and diagnostics

For each saved time point, the simulations stored

$$\phi_{\mathrm{Dps}}(\mathbf{r}), \qquad \phi_{\mathrm{DNA}}(\mathbf{r}), \qquad \phi_{\mathrm{Solv}}(\mathbf{r}), \qquad \phi_T(\mathbf{r}) = \phi_{\mathrm{Dps}}(\mathbf{r}) + \phi_{\mathrm{DNA}}(\mathbf{r}).$$

The following consistency checks were performed:

$$\langle \phi_{\mathrm{Dps}} \rangle = \bar{\phi}_{\mathrm{Dps}}, \tag{70}$$

$$\langle \phi_{\mathrm{DNA}} \rangle = \bar{\phi}_{\mathrm{DNA}}, \tag{71}$$

$$\phi_{\mathrm{Dps}} + \phi_{\mathrm{DNA}} + \phi_{\mathrm{Solv}} = 1, \tag{72}$$

$$\phi_T = \phi_{\mathrm{Dps}} + \phi_{\mathrm{DNA}}. \tag{73}$$

Thus Dps and DNA are individually conserved, the solvent follows from incompressibility, and the dense co-condensate field is recovered by summing the two solute fields.

#### 9 Compactness of DNA due to Dps

Beyond single-particle motion tracking of the Dps bead, the collapse of the DNA chains themselves is slow and heterogeneous. Thus, we require a measurement that describes how the overall polymer changes as Dps binding drives condensation. The radius of gyration, $R_g$, provides this information by measuring how the DNA mass is distributed about its center of mass. Smaller $R_g$ values indicate a tighter compaction, while larger values correspond to more extended configurations. Since our simulations are quasi-2D, we computed $R_g$ using the $x$ and $y$ coordinates of DNA bead positions.

Computing $R_g$ begins with finding the gyration tensor, which characterizes the spatial distribution of DNA beads relative to the DNA chain's center of mass. Bead coordinates were centered relative to the chain's center of mass, $(x_{CM}, y_{CM})$, and used to form the tensor.

$$G = \begin{pmatrix} G_{xx} & G_{xy} \\ G_{xy} & G_{yy} \end{pmatrix} \tag{74}$$

where $G_{xx} = \frac{1}{N}\sum_{i=1}^{N}(x_i - x_{CM})^2$, $G_{yy} = \frac{1}{N}\sum_{i=1}^{N}(y_i - y_{CM})^2$, and $G_{xy} = \frac{1}{N}\sum_{i=1}^{N}(x_i - x_{CM})(y_i - y_{CM})$. Here, $x_i$ and $y_i$ are the bead positions of the DNA, and $N$ is the number of beads in the DNA chain. The eigenvalues of the gyration tensor represent the principal spatial extents of the polymer along its principal axes. Thus, the sum of the eigenvalues measures the total spatial spread of the chain, which yields the squared radius of gyration. Since our simulations are carried out in a quasi-2D box ($L_z \ll L_x = L_y$), the eigenvalue corresponding to the $z$-direction is assumed to be zero. We diagonalize $G$ to obtain its two non-zero eigenvalues $\lambda_1$ and $\lambda_2$, and then compute $R_g^2 = \lambda_1 + \lambda_2$, which yields $R_g = \sqrt{\lambda_1 + \lambda_2}$. We perform this calculation for simulations where $\phi_{\text{DNA}} = 0.0013, 0.0026$ and $\phi_{\text{Dps}} = 0.0236, 0.0707$. Additionally, we have kept the interaction strengths fixed, where $\varepsilon_{\text{Dps:DNA}} = 1.5$ and $\varepsilon_{\text{Dps:Dps}} = \varepsilon_{\text{DNA:DNA}} = 1.0$.

Computing the radius of gyration in this manner provides information on how one entire chain of DNA behaves under compaction/binding by Dps. While this may provide information on how a DNA chain compacts from Dps globally, it can obscure localized behavior because different regions of the DNA chain do not collapse simultaneously. Dps beads in the system can bind at different locations along the DNA chain, which can produce heterogeneous domains that compact on different timescales. To resolve this behavior, each 500-bead DNA chain was divided into five contiguous 100-bead segments, and $R_g$ was computed for each segmented chain. This segmentation reduces the influence of large-scale translations and bending of the entire DNA chain while providing a local measure of compaction that is more sensitive to where condensation first develops. Choosing substantially shorter segments of DNA would introduce excessive statistical fluctuations because there would not be enough beads to contribute to the computation of $R_g$; and choosing larger segments would reintroduce the behavior we would see from computing $R_g$ globally over the entire chain. Thus, a segment length of 100-beads provides a balance between spatial resolution and statistical robustness.

We compute the radius of gyration as a function of time. For each simulation seed, the radius of gyration was computed independently for each 100-bead DNA segment in a chain. These segment-level measurements were then averaged across chains and simulation seeds to obtain an ensemble-averaged $R_g(t)$ curve for each DNA and Dps concentration. Because segments from the same DNA chain are not fully independent, the segmented $R_g$ values were used to resolve local compaction rather than to artificially increase the number of independent samples. The standard error of the mean was therefore interpreted as the uncertainty in the averaged compaction trajectory, while the stretched-exponential fit was applied to the ensemble-averaged $R_g(t)$ curve from all four seeds.

To quantify the kinetics of this relaxation, $\overline{R_g(t)}$ data was fit with a stretched exponential

$$R_g(t) = R_g^{\infty} + (R_g^0 - R_g^{\infty}) \exp\left[-(t/\tau)^{\beta}\right]. \tag{75}$$

$R_g^{\infty}$ is the steady state value that $R_g(t)$ approaches over the course of the simulation and describes how compact the final DNA configuration becomes. The parameter $\tau$ describes how fast DNA compaction occurs, and $\beta$ denotes the stretching exponent relating to the range of the distribution of relaxation times. This stretched exponential captures both the rapid initial decrease and the slower approach to the dynamic, arrested state.

This local analysis is particularly important because the DNA does not collapse uniformly. Instead, compaction initiates in localized regions where Dps first accumulates and subsequently propagates along the chain. Segmenting the polymer therefore allows heterogeneous relaxation dynamics to be resolved that would otherwise be masked by a single global radius of gyration.
